# NK cells control the progression of myelodysplastic syndrome but become initial disease target in *NUP98-HOXD13* mouse model

**DOI:** 10.1101/2024.12.11.627924

**Authors:** Gladys Telliam-Dushime, Maciej Ciesla, Henrik Lilljebjörn, Jonas Ungerbäck, Ouyang Yuan, Dang Nghiem Vo, Olga Kotova, Thoas Fioretos, David Bryder, Cristian Bellodi, Ewa Sitnicka

**Affiliations:** Division of Molecular Hematology, Department of Laboratory Medicine, Lund Stem Cell Center, Lund University, Sweden; Division of Clinical Genetics, Department of Laboratory Medicine, Lund University, Sweden

**Keywords:** NK cell defects, myeloid malignancies, NK cell development and homeostatic proliferation

## Abstract

Studies in *NUP98/HOXD13* mouse model (NHD13^tg^), progressing from myelodysplastic syndrome (MDS) to different forms of leukemia, demonstrated that T cells had a limited anti-leukemia effect, suggesting the involvement of other immune cells. Natural killer (NK) cells control viral infection and cancer. In MDS and acute myeloid leukemia (AML), patients often acquire disease-induced NK cell dysfunctions. Here, we report that NK cells from NHD13^tg^ mice were reduced before the MDS-onset and specific NK cell depletion accelerated the disease progression and severity. NK cells from NHD13^tg^ mice showed perturbed differentiation and impaired IL-15/IL-2 responses. These defects were cell-intrinsic and mainly affected the KLRG1^+^ mature NK cells. The expression of Nfil3, Klf2 and Id2 genes, crucial for NK cell development, homeostasis and IL-15 responsiveness, was altered in immature NK cells from NHD13^tg^ mice. Interestingly, these genes were changed in MDS and AML bone marrow patient-samples compared to healthy donors. Our findings highlight a critical role for NK cells in controlling MDS progression and identify new genetic markers for MDS and AML.

## INTRODUCTION

Myelodysplastic syndrome (MDS) and acute myeloid leukemia (AML) are clonal disorders characterized by the accumulation of genetic aberrations in hematopoietic stem and progenitor cells (HSPCs) that leads to myeloid progenitors (blasts) expansion in the bone marrow (BM) and periphery (*1, 2*).

MDS and AML are two of the most common myeloid malignancies, associated with poor prognosis (*3*), and 30% of MDS patients progresses to AML.

Natural killer (NK) cells are key components of the innate immune system directly involved in tumor immune surveillance (*4*) as demonstrated in multiple mouse studies (*5*). In humans, an increased risk of cancer has been linked to NK cell-deficiencies (*6*), and NK cells from patients with MDS and AML have poor cytotoxic functions and reduced expression of specific cell surface cytotoxic receptors (*7, 8*). The severity of NK cells impairment correlates with more advanced cancer stages (*9*).

NUP98 is a component of the nuclear pore complex responsible for bidirectional trafficking to the nucleus (*10*). NUP98 protein level affects expression of genes involved in cell cycle and differentiation (*10*). Many fusions involving *NUP98* gene have an impact on hematopoietic precursors differentiation process (*10*). The *NUP98-HOXD13* (NHD13) fusion gene is found in patients with myeloid malignancies and acts as an aberrant transcription factor that leads to leukemic transformation (*10*) by inducing the expression of genes frequently involved in MDS pathogenesis such as *HOXA 7, 9 or 10* (*11*).

Mice expressing *NUP98/HOXD13* transgene (NHD13^tg^ mouse model) were previously generated and characterized (*12*). NHD13^tg^ mice develop MDS between 4 to 7 months of age, which can progress to different forms of leukemia including AML and T-cell lymphoblastic leukemia (T-ALL) around 10 to 14 months (*12*) but, for most cases, NHD13^tg^ mice die from severe forms of MDS (*12*). NHD13^tg^ mice progressively develop a peripheral blood cytopenia and leukopenia, a defect in lymphoid B and T cells, and an increase of immature myeloid cells (*13, 14*), which can be transferred upon transplantation (*13, 14*).

Crossing NHD13^tg^ mice with RAG-1 knockout mice led to only modest decrease in survival and slightly more frequent progression to leukemia, suggesting a mild anti-leukemia effect of T-lymphocytes and a potential involvement of other immune cells (*15*).

Although NK cells represent the main cell type responsible for cancer immune-surveillance, their development, function and involvement in the disease progression have never been investigated in NHD13^tg^ mice.

Here we show that specific NK cell depletion leads to striking acceleration of disease progression from MDS to leukemia and overall decreased survival. In NHD13^tg^ mice, the pool of mature NK cells is significantly impaired already at the pre-MDS blast-free stage. Multiple NK cell defects; reduced cytokine responsiveness, homeostatic proliferation and induced transcriptional changes; together contributed to the progressive exhaustion of the NK cell compartment. Finally, we identified potential novel genetic diagnostic markers for MDS and AML. All together our study highlights the direct and critical role of NK cells in controlling disease evolution from MDS to leukemia.

## RESULTS

### NK cell defects occur before the MDS onset and progress with the disease advancement in NHD13^tg^ mouse model

First, we investigated the distribution of different blood lineages in 6-9 week-old pre-MDS blast-free stage and 5-7 month-old animals with an established MDS-like phenotype. In agreement with previous studies (*12, 14*), at the MDS stage, the frequency of B and T cells was significantly reduced, the proportion of myeloid cells (CD11b^+^Gr1^+^) was significantly increased, and myeloblast cells (CD11b^+^cKIT^+^) which also expressed low level of Gr1 (data not shown) were clearly detectable in the spleens of NHD13^tg^ mice compared to WT littermate controls (Figure 1A-B) with an expected strong correlation between myeloblasts and lymphocytes proportion (Figure 1C). Surprisingly, we found that both the proportion and total number of peripheral mature NK cells (mNK) (Ter119^-^CD19^-^CD3^-^NK1.1^+^) were significantly reduced in the PB and spleen already in pre-MDS 6-9 week-old NHD13^tg^ mice (Figure 2A-C). The disease progression to the MDS stage in 5-7 month-old NHD13^tg^ mice was associated with further NK cell depletion, both in the PB and spleen (Figure 2A-C). The total WBC counts were also significantly lower in the spleens of NHD13^tg^ mice (Supplementary Figure 1A) but not in the PB (Supplementary Figure 1B) at both disease stages. Importantly, NK cells from NHD13^tg^ mice were carrying the *NUP98-HOXD13* fusion gene (Figure 2D). Interestingly, in 5-7 month-old NHD13^tg^ mice with detectable NK cells (Figure 2D), an enhanced NK cell reduction (median NK cell frequency ≤ 0.96%) correlated with more severe MDS-like disease phenotypes, including clear association between low NK cell proportion, elevated level of myeloid cells, increased proportion of blasts (cKIT^+^) and reduction of B and T cells (Figure 2E-F).

**Figure 1.**
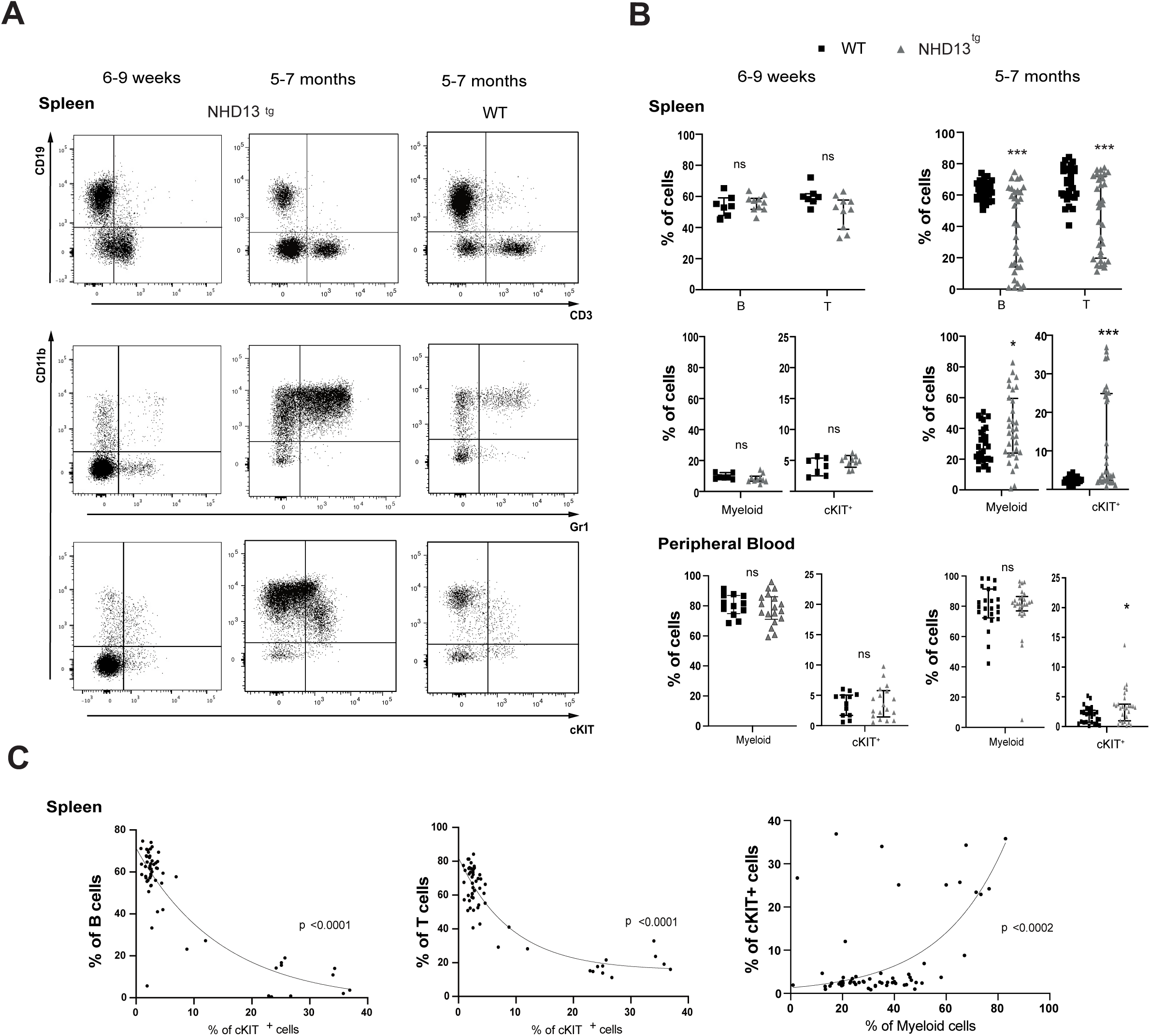
Cell distribution in the spleen of NHD13^tg^ mice at pre-MDS and MDS stages. **(A)** Representative FACS^TM^ plots of B (CD19^+^) and T (CD3^+)^ lymphocytes, myeloid (CD11b^+^Gr1^+^) and blast (Gr1^+^cKIT^+^) cells within Ter119^-^ Live splenocytes from 6-9 weeks old (left panels), 5-7 months old NHD13^tg^ mice (middle panels) and 5-7 months old WT littermate controls (right panel). All negative values indicate a marker that was gated out, therefore each new gate represents 100% of remaining cells. **(B)** Percentage of T and T lymphocytes, myeloid and blast cells in the spleen from 6-9 weeks (left panels) and 5-7 months old (right panels) NHD13^tg^ mice (grey triangels) and WT littermate controls (black square). At least 7 mice were analyzed per group in 3-4 independent experiments. Statistical differences between the groups were determined using unpaired Mann-Whitney U test *p<0.05, ***p<0.001, ns=not significant. **(C)** Representative correlation plots between the proportion of blast (cKIT^+^) cells and the proportion of B, T and myeloid cells from the 5-7months old NHD13^tg^ and WT littermate control mice. The p values are indicated within each plot.

**Figure 2.**
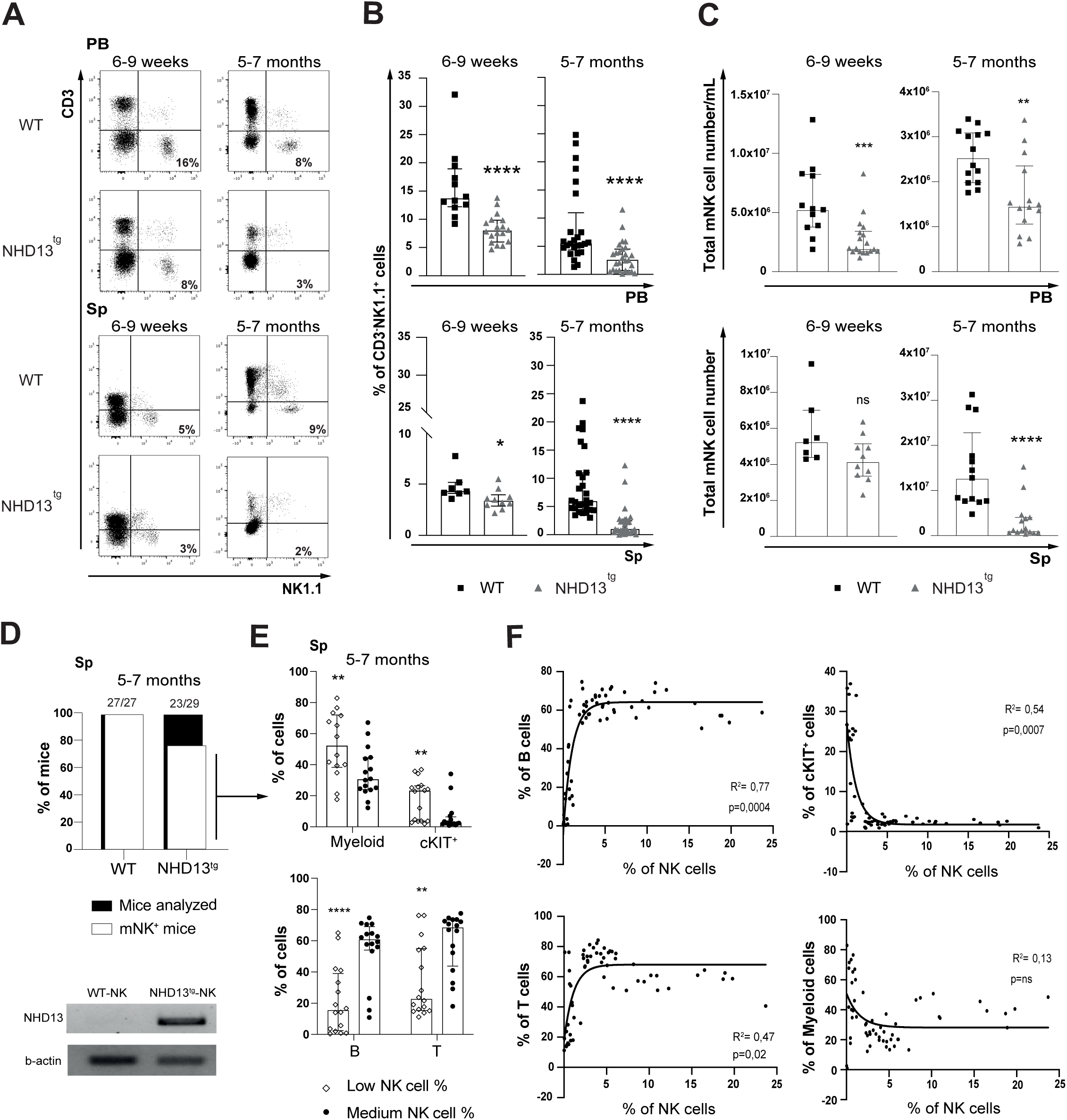
Reduction in mature NK cells occurs already at the pre-MDS stage and progresses with the disease advancement to MDS in NHD13^tg^ mice. **(A)** Representative FACS™ plots of peripheral blood (PB) (top) and splenic (Sp) (bottom) mature NK (mNK) cells (Ter119^-^CD19^-^CD3e^-^NK1.1^+^) from NHD13^tg^ and littermate control 6-9 weeks (left) and 5-7 months (right) old mice. Numbers in each quadrant within NK cell gate indicate mean values corresponding to each animal group. **(B)** Percentage of mNK cells in PB (top) and spleen (bottom) in 6-9 weeks (left) and 5-7 months (right) old WT (black squares) and NHD1 3^tg^ (grey triangles) mice. Each bar represents the median with interquartile range. A minimum of 7 mice was analyzed per group in 3-4 independent experiments. **(C)** Total number of mNK cells calculated based on the total WBC (PB) (top) and spleen (Sp) (bottom) cell counts. **(D)** Histogramme representing the proportion of mice analyzed with detectable mNK cells. For each group mNK cells (Lin^-^CD3^-^CD122^+^NK1.1^+^DX5^+^CD11b^+^) were sorted and the expression of *NUP98-HOXD13* fusion gene together with the expression of beta-actin was detected by RT-PCR. **(E)** Frequencies of myeloid cells (Ter119^-^ CD19^-^CD3^-^NK1.1^-^CD11b^+^Gr1^+^), blast cells (Ter119^-^cKIT^+^), B (Ter119^-^CD19^+^) and T (Ter119^-^CD19^-^CD3^+^NK1.1^-^) cells from 5-7 months old NHD13^tg^ mice with low (open circle) or medium (black circle) levels of mNK cells based on the median value of 0,96%. **(F)** Representative correlation plots between the NK cell proportion and the proportion of B, T, cKIT^+^ and myeloid cells from 5-7 months old NHD13^tg^ and WT littermate cohort. The R squared values and p values are indicated within each plot. Statistical differences between groups were determined using unpaired Mann-Whitney U test *p ≤0.05, **p≤0.01, ***p ≤0.001, ****p ≤0.0001, ns= non significant.

These results support that already in young MDS-free NHD13^tg^ mice, the pool of NK cells is reduced and becomes increasingly defective with the disease progression correlating with the severity of the MDS phenotype.

### NK cell development in NHD13^tg^ mice is delayed at the M1 to M2 transition stage and the mature KLRG1^+^ NK cell population is selectively lost

The current model of mouse NK cell development comprises multiple cellular stages (*16*). From the NKP progenitor, differentiating NK cells acquire the expression of NK1.1 receptor, then upregulate DX5, CD11b (Mac1) and CD27 molecules. CD27 and CD11b expression separates mNK cells (NK1.1^+^DX5^+^) into three sequential stages: M1 (CD27^+^CD11b^-^), M2 (CD27^+^CD11b^+^) and M3 (CD27^-^CD11b^+^) (*17*). In addition, the presence of MHC class I inhibitory killer cell lectin-like receptor G1 (KLRG1) allows further characterization of NK cell maturation status (*18*) (Supplementary Figure 2A).

To establish at which developmental stage the NK cell defect occurs in NHD13^tg^ model, we investigated BM and splenic conventional NK cell subsets, and their progenitors in 6-9 week-old pre-MDS and 5-7 month-old MDS NHD13^tg^ mice. Interestingly, we found in the spleens of pre-MDS NHD13^tg^ mice, an accumulation of M1 mNK cells (CD27^+^CD11b^-^KLRG1^-^) (Figure 3A-B, Supplementary Figure 2B) and a reduction of M2 mNK cells (Lin^-^ CD122^+^NK1.1^+^DX5^+^CD27^+^CD11b^+^) (Figure 3A-B, Supplementary Figure 2B) that together suggests a potential developmental defect at the transition from the M1 to M2 stage.

**Figure 3.**
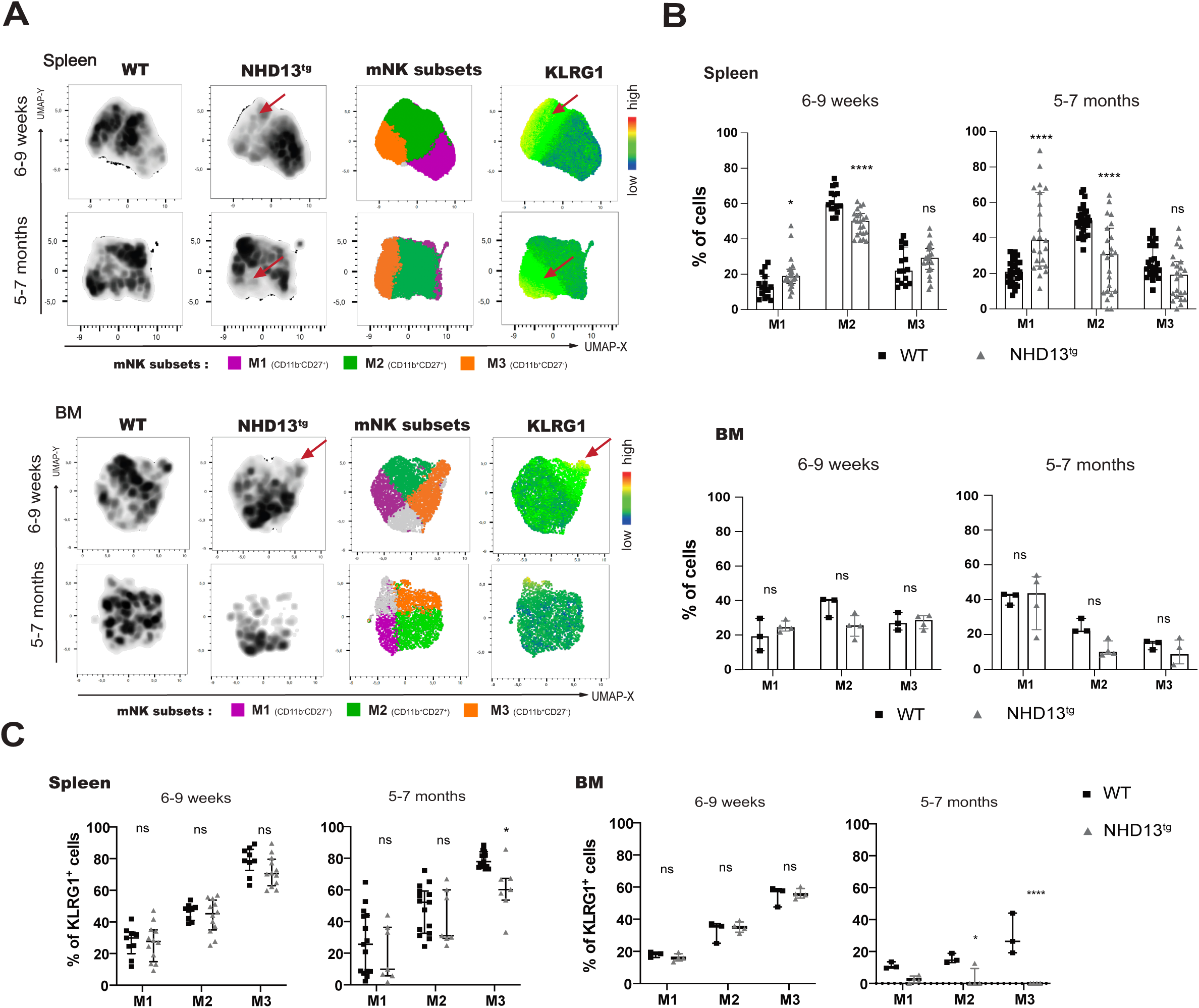
NK cell differentiation in NHD13^tg^ mice. **(A)** Multidimensional flow cytometry using Uniform Manifold Approximation and Projection (UMAP) density map generated from spleen (top panel) and BM cells (bottom panel) from 6-9 weeks and 5-7 months old NHD13^tg^ mice and WT littermate controls. Mature NK (mNK) cell subsets localization is indicated within the maps. Heatmap statistics shows the expression level of KLRG1, a key surface marker of mature and functional NK cells. Arrows indicate cells expressing high levels of KLRG1. All maps were generated from FlowJo™ UMAP plugin after concatenation of minimum 3 mice per group. **(B)** Percentage of mNK cells in the spleen (top panels) and BM (bottom panels) in 6-9 weeks and 5-7 months old NHD13^tg^ mice (grey triangles) and WT littermate controls (black squares). **(C)** Percentage of KLRG1^+^ cells within mature NK cell subsets in the spleen (left panels) and BM (right panels) in 6-9 weeks and 5-7 months old NHD13^tg^ mice (grey triangles) and WT littermate controls (black squares). Minimum three to eighteen mice were analyzed per group. Statistical differences were determined using Two-Way ANOVA multiple comparisons *p ≤ 0.05, **p≤0.01, ****p ≤ 0.0001, ns= not significant.

During the disease progression to the MDS stage, approximately 20% of 5-7 month-old NHD13^tg^ mice had no detectable levels of mNK cells in the spleen (Figure 2D). The fraction of 5-7 month-old NHD13^tg^ mice with detectable splenic mNK cells showed consistent accumulation of M1 mNK cells (Lin^-^CD122^+^NK1.1^+^DX5^+^CD27^+^CD11b^-^), decrease in M2 mNK cells (Lin^-^CD122^+^NK1.1^+^DX5^+^CD27^+^CD11b^+^) and selective loss of mNK cells (CD11b^+^KLRG1^high^) also found in the BM, compared to the WT littermate controls (Figure 3A-C, Supplementary Figure 2B). The spleen cellularity was significantly reduced in NHD13^tg^ mice at both disease-stages compared to WT controls, further contributing to an overall reduction in the total mNK cell number (Figure 2C, Supplementary Figure 1A, Supplementary Figure 2).

Detailed analysis of BM from pre-MDS mice revealed a depletion of the lympho-myeloid (CD27^+^cKIT^+^CD135^+^) cell cluster from the t-SNE map (Supplementary Figure 3); while the proportion of CLPs (CD244^+^CD27^+^CD127^+/-^cKIT^+^CD135^+^CD122^-^) was significantly reduced, the pre-NKP cells (CD135^-^CD122^-^) were increased. However, the r-NKP progenitor pool (CD135^-^CD122^+^) remained unaltered compared to the WT littermate counterpart, suggesting that changes within the early BM progenitors do not contribute to the early pre-MDS reduction of conventional mNK cell pool (Supplementary Figure 3).

Together these results suggest that the NK cell loss in NHD13^tg^ mice is caused by a perturbed transition from the M1 to M2 developmental stages occurring at the pre-MDS stage, that specifically affects KLRG1^high^ mNK cells.

### NK cell defect in NHD13^tg^ mice is cell intrinsic

To determine whether the identified NK cell defects are cell intrinsic, we performed competitive transplantations using unfractionated BM cells from 8 week-old pre-MDS NHD13^tg^ CD45.2 mice or WT littermate controls together with competitor BM cells from WT B6SJL CD45.1 mice. At 4 months after transplantation, donor-derived reconstitution of the NK cell compartment in PB was significantly lower in recipient mice transplanted with cells from NHD13^tg^ donors compared to those transplanted with WT cells (Figure 4A, Supplementary Figure 4), whereas the lineage reconstitution from competitor cells in both chimera groups was not altered (Supplementary Figure 4B). At 6 months post-transplantation, NK cell generation from NHD13^tg^ donors was further impaired and accompanied by a significant increase in myeloid cells and blast cells (cKIT^+^) (Figure 4A), reproducing the NK cell defect and MDS-like phenotype found in NHD13^tg^ donor mice. The final analysis 8 months post-transplantation gave the same results including significant lymphopenia (Figure 4A). Whereas all recipient mice transplanted with CD45.2 BM cells from control mice survived, the survival probability of animals transplanted with NHD13^tg^ BM cells was significantly lower (Figure 4B). Importantly, the analysis of lineage reconstitution in the spleens revealed that mNK cell generation from transplanted BM cells from NHD13^tg^ mice was also affected leading to the accumulation of M1 CD27^+^ mNK subset and decrease of KLRG1^+^ mNK cells, as observed in NHD13^tg^ donor mice (Figure 4C-D). The UMAP analysis of mNK cells generated from the transplanted competitor CD45.1 BM cells showed no significant changes in immature M1 CD27^+^ cells and KLRG1^+^ mNK cell proportion (Figure 4C-D).

**Figure 4:**
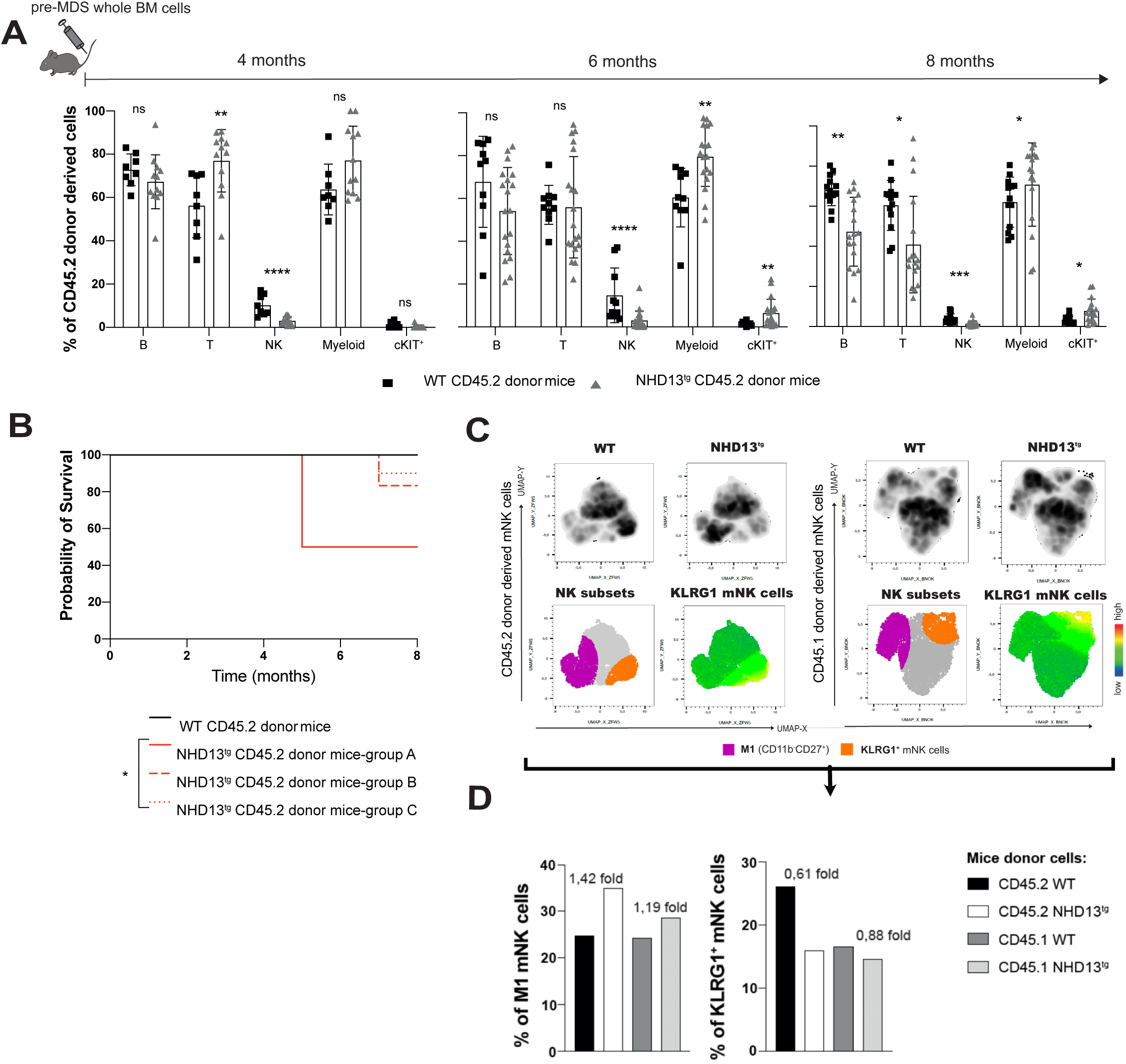
Blood lineage reconstitution in chimera mice after competitive transplantation. **(A)** Percentage of CD45.2 donor-derived peripheral blood cell types at 4, 6 and 8 months post-transplantation with CD45.2 NHD13^tg^ (grey triangles) or WT littermates (blacks quares) donor cells together with CD45.1 WT competitor cells from B6SJL mice. Data are from 3 independent experiments with minimum 8 recipient mice transplanted per group. Statistical differences were determined using 2-way ANOVA with multiple comparison ** p≤0.01, **** p≤0.0001, ns=not significant. **(B)** Kaplan Meier analysis of recipient mice from 3 independent experiments presented individually (groups A, B and C) transplanted with CD45.2 NHD13^tg^ or CD45.2 WT littermate donor cells showing the probability of survival. The statistical difference was determined using Gehan-Breslow-Wilcoxon test (p=0.0131) * p≤ 0.05. **(C)** Multidimensional flow cytometry with Uniform Manifold Approximation and Projection (UMAP) density map of CD45.1 or CD45.2 donor-derived mNK cell subsets in the spleens from recipient mice analyzed at 8 months post transplantation. All maps were generated from FlowJo™ UMAP plugin after concatenation of minimum 5 mice per group. M1 mature NK cell subset and KLRG1^+^ mNK cells localization are indicated within the map. Heatmap statistics shows the expression level of KLRG1, a key surface marker of mature and functional NK cells. **(D)** Percentage of CD45.1 or CD45.2 donor derived M1 mature NK cells and KLRG1^+^ cells within the UMAPs. Numbers above histograms indicate the mean fold change in M1 mNK cell subsets and KLRG1 expressing cells generated according to each respective reference CD45.1 cells.

Together these data support that NK cell defects in NHD13^tg^ mice are cell intrinsic, that is further supported by the detection of *NUP98-HOXD13* fusion gene in NK cells purified from NHD13^tg^ mice (Figure 2D) significantly affecting their survival probability.

### Selective NK cell depletion accelerates disease progression to leukemia

Next, we selectively depleted the mNK cell compartment in NHD13^tg^ mice at the MDS stage using the anti-NK1.1 antibody treatment (*19*). The loss of NK cell pool led to a critical decrease in the animal survival compared to the isotype control-injected group (Figure 5A-B). The NK cell-depleted NHD13^tg^ mice were showing clear signs of disease acceleration and progression towards leukemia already within 3 months post-injection which manifested as an increased myeloid cell proportion, the presence of 10 to 30% of blast cells in the PB (in 2 cases were lymphoblasts with 5 to 16% of Gr1^-^cKIT^+^ cells and a doubled size of the T cell pool) (Figure 5C, Supplementary Figure 5), and severe lymphopenia (in some cases) (Supplementary Figure 5). The elevated levels of blast and myeloid cells in NHD13^tg^ mice treated with anti-NK1.1 antibody were comparable to those observed in NHD13^tg^ mouse at myeloid leukemia stage (non-treated, positive reference control) (Figure 5C-D). The group of NHD13^tg^ mice injected with control antibody, showed minor or no signs of disease progression, as the levels of myeloid and blast cells remained the same after the treatment (except for one case reaching 5% of blast) (Figure 5C-D).

**Figure 5.**
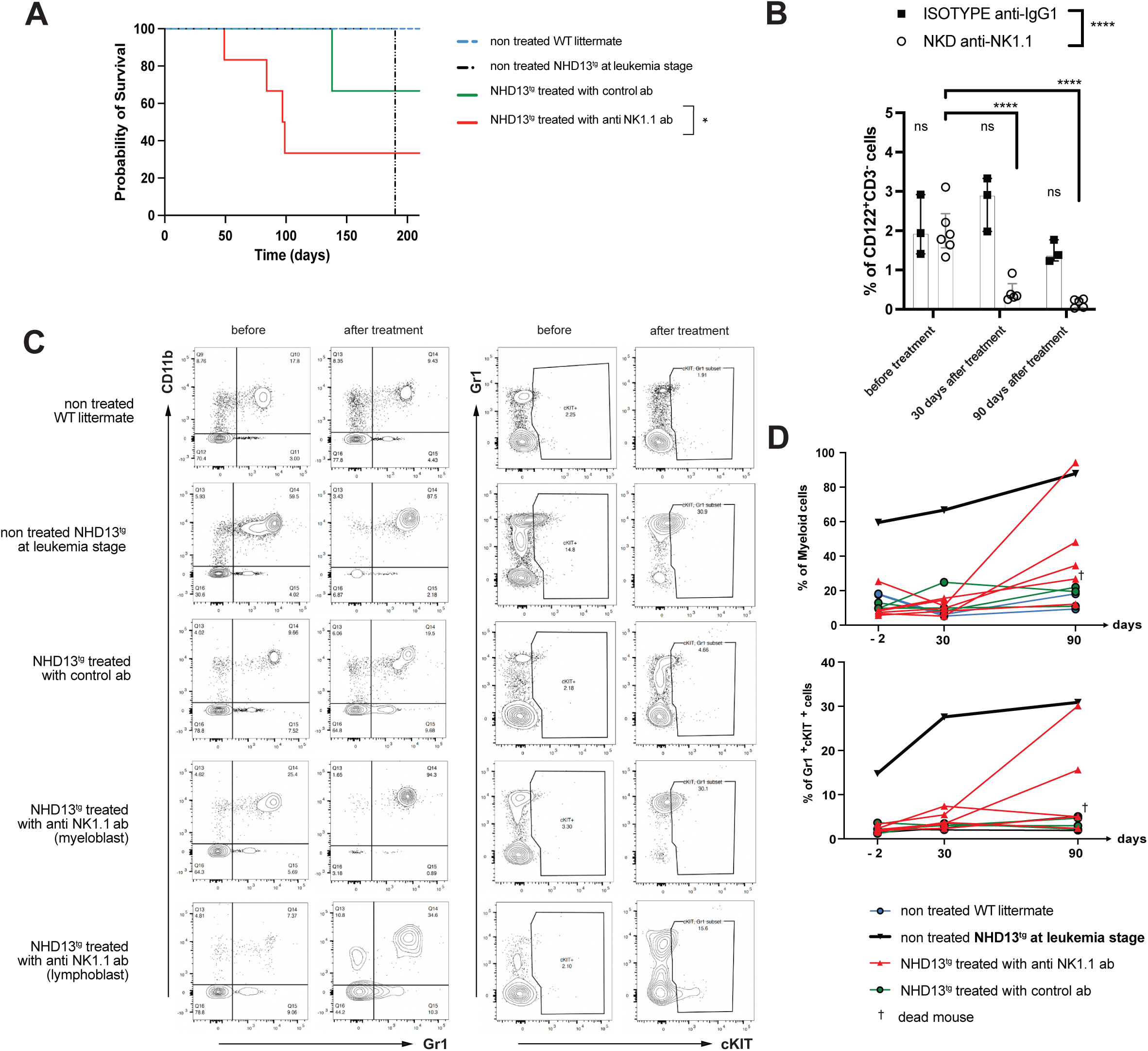
Selective NK cell depletion leads to accelerated disease progression. **(A)** Kaplan Meier analysis of NHD13^tg^ mice cohorts injected with specific anti-NK1.1 (n = 6) or isotype control antibody (n = 4). **(B)** Summary histogram of NK cell percentage (CD122^+^CD3^-^) before and 1 or 3 months after treatment with anti-NK1.1 antibody or isotype control antibody. **(C)** FACS™ profile of myeloid cells (CD11b^+^Gr1^+^) and blast cells (Gr1^+^cKIT^+^ or Gr1^-^ cKIT^+^) before and 3 months after treatment. **(D)** Percentage of myeloid cells (CD11b^+^Gr1^+^), blast cells (Gr1^+^cKIT^+^) before (-2 days) and after (30 and 90 days) specific NK cell depletion. A cohort of 10 to 11 months old NHD13^tg^ mice injected with isotype control antibody was used as a negative control. Age-matched WT littermate mice were used as a negative reference, and a 1-year old NHD13^tg^ mouse at myeloid leukemia stage was used as a positive reference. N/A= non applicable, † = mouse dead at the indicated time point.

These results clearly demonstrate that specific NK cell loss in NHD13^tg^ mice led to an accelerated disease, supporting the direct role for NK cells in controlling disease progression.

### Peripheral mNK cells in pre-MDS NHD13^tg^ mice show alteration in the responsiveness to IL-15 and IL-2

Based on our finding that M1 mNK cells accumulation coinsides with significant reduction of the M2 subset in pre-MDS NHD13^tg^ mice (Figure 3, Supplementary Figure 2), we hypothesized that the impaired response to cytokines may affect mature NK cell homeostasis. To investigated the responsiveness to key regulatory cytokines controlling NK cell development, proliferation and function; we evaluated NK cell apoptosis, survival and proliferation by culturing in competetion equal numbers of splenocytes from NHD13^tg^ and WT littermate control mice pre-labelled with different cell-tracing dyes Cell-Trace Far-Red (CTR) and Cell-trace Violet (CTV) respectively, in an increased concentrations of IL-15 or IL-2. In the presence of IL-15, approximately 75% of mNK cells maintained in culture for 7 days belonged to CTV-stained WT littermate counterpart (Figure 6A-B). Interestingly, the highest IL-15 concentration (40ng/ml) could partially support survival of CTR-labelled NK cells from NHD13^tg^ mice (Figure 6B). Competitive culture in IL-2 consistently led to the survival of 75 to 100% of WT CTV-stained NK cells, regardless of the IL-2 concentrations (Figure 6C-D) with only few CTR-labelled NK cells from NHD13^tg^ mice surviving (Figure 6C-D). There was no increase in early apoptosis within mNK cells from NHD13^tg^ compared to WT littermate controls (Supplementary Figure 6A).

**Figure 6.**
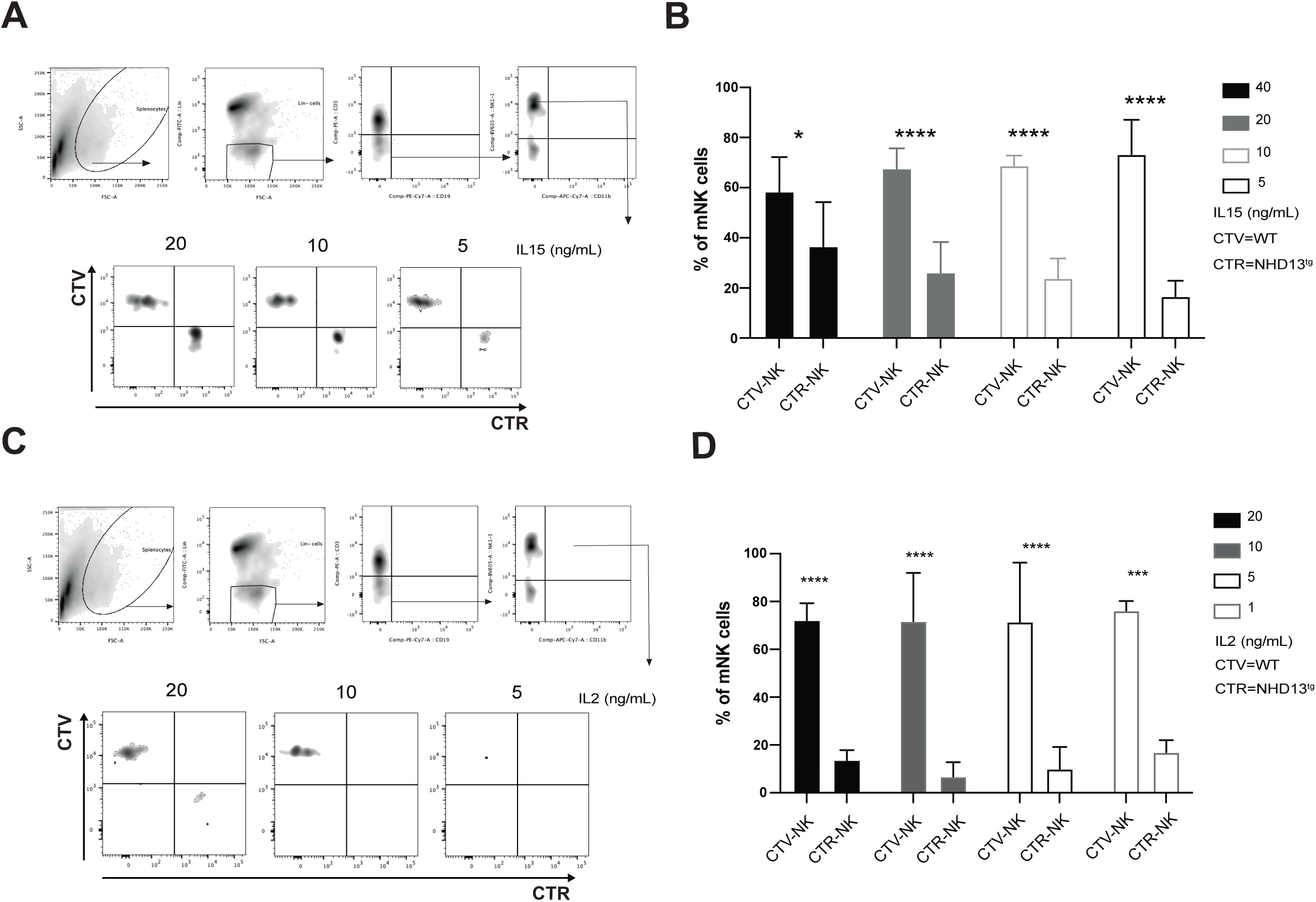
Conventional mature NK cells in NHD13tg mice have an impairment in the response to IL-15 and IL-2 signaling. **(A)** Gating strategy and representative FACS™ plots of splenocytes isolated from 6-9 weeks old NHD13^tg^ mice or WT littermates stained with cell trace dyes, Cell Trace Far-Red (CTR) and Cell Trace Violet (CTV), respective.ly. Labelled cells were cultured at a ratio 1:1 in competition for 7 days in presence of increased concentration of IL-15. The CTV against CTR fluorescence is displayed within mNK cell (Lin^-^CD3^-^CD19^-^ NK1.1^+^) gate. **(B)** Percentage of mNK cells expressing CTR (NHD13^tg^-derived) or CTV (WT littermate-derived) in response to different IL-15 concentrations (40, 20, 10 and 5 ng/ml depicted as black, grey and white bars, respectively). **(C)** Gating strategy and representative FACS™ plots of splenocytes from 6-9 weeks old NHD13^tg^ mice or WT littermates stained with cell trace dyes, Cell Trace Far-Red (CTR) and Cell Trace Violet (CTV), respectively. Labelled cells were cultured in competition for 7 days in presence of increased concentration of IL-2. The CTV against CTR fluorescence is displayed within mNK cell (Lin^-^CD3^-^CD19^-^NK1.1^+^) gate. **(D)** Percentage of mNK cells expressing CTR (NHD13^tg^-derived) or CTV (WT littermate-derived) in response to different IL-2 concentrations (20, 10, 5 and 1 ng/ml depicted as black, grey and white bars, respectively). Each bar represents the mean value with SD. At least 5 mice were analyzed per group in 2 independent experiments. Statistical differences between groups were determined using ANOVA with multiple comparison *p ≤ 0.05, ***p≤0.001, ****p≤0.0001.

Next, we analyzed the relative expression level of gene transcripts required for IL-15 and IL-2 signaling (Supplementary Figure 6B). All mRNA were expressed at similar levels in CD11b^+^ M2 mNK cell subset sorted from NHD13^tg^ and WT littermate mice. Interestingly, while the expression of Stat5a/b was higher in CD11b^-^ M1 mNK cells from NHD13^tg^ mice, there was no difference in CD122 levels between both genotypes (Supplementary Figure 6B). Therefore, the failure of NHD13^tg^ NK cells to respond and compete for IL-15 and IL-2 signals was not due to a lack of downstream signaling partners nor due to an altered receptor sequence (data not shown). We found no evidence for a lack of survival signal, neither an increased level of pro-apoptotic signal, as relative expression levels of several key Bcl2 family members were not significantly altered (Supplementary Figure 6C-D).

Despite the impaired IL-15 responsiveness, mNK cells from NHD13^tg^ mice maintained their functional properties including cytotoxic activity: CD107a up-regulation (*20*), and the ability to release IFN-γ in response to IL-12/IL-18 stimulation. The expression level of key cell-surface receptors and genes involved in NK cell cytotoxicity was comparable in NHD13^tg^ and WT mice (Supplementary Figure 7).

Collectively, these results suggest that the reduced size of the mNK cell pool at the pre-MDS stage is caused by an impaired response to IL-15 signaling, leading to the loss of KLRG1^+^ mNK cells (Figure 3B) that require IL-15 for their homeostatic proliferation.

### M1 NK cells from NHD13^tg^ mice show pre-mature transcriptional profile compatible with the M2 developmental stage

To explore molecular mechanisms responsible for the NK cell defects observed in NHD13^tg^ mice at the pre-MDS stage, we compared the gene expression profiles of M1, M2 and M3 conventional mNK cell subsets FACS sorted from 6-9 week-old NHD13^tg^ and their WT littermate counterparts (Figure 7, Supplementary Figure 8). The transcripts encoding genes relevant to the accuracy and quality of the experimental procedures including: Il2rß, Ncr1 and Eomes were expressed in all mNK cell subsets sorted from both genotypes (Figure 7A-C). The expression levels of cell surface markers including CD11b, CD27, CD2, CD62L, CD96 and Cxcr1 related to differentiation and function were compatible with the NK cell maturation stages (Figure 7A-B, Supplementary Figure 8A). Finally, Hoxa9, a known target of *NUP98- HOXD13* fusion gene was only detectable in mNK cell subsets sorted from NHD13^tg^ mice (Figure 7A-B). Interestingly, as illustrated in the Euclidean Distance analysis of different NK cell subsets, based on their gene expression pattern, M1 cells from NHD13^tg^ mice clustered closely to M2 cells from WT littermate control (Figure 7E) highlighting the advanced maturation in M1 cells from NHD13^tg^ mice compared to their WT counterpart (Figure 7E). Specifically, transcription factors Klf2, Tox, Tcf7, Gata3, Irf2, Nfil3 and T-bet, all known to regulate NK cell maturation (*21, 22*) were differentially expressed in M1 cells from NHD13^tg^ mice and resembled M2 cells from WT littermate mice (Figure 7A-C, Supplementary Figure 8B). The transcription factor Id2, previously reported to control NK cell development and IL- 15 responsiveness (*23, 24*) was expressed at reduced levels in all mNK cell subsets from NHD13^tg^ mice compared to their respective WT counterparts (Figure 7A-C).

**Figure 7.**
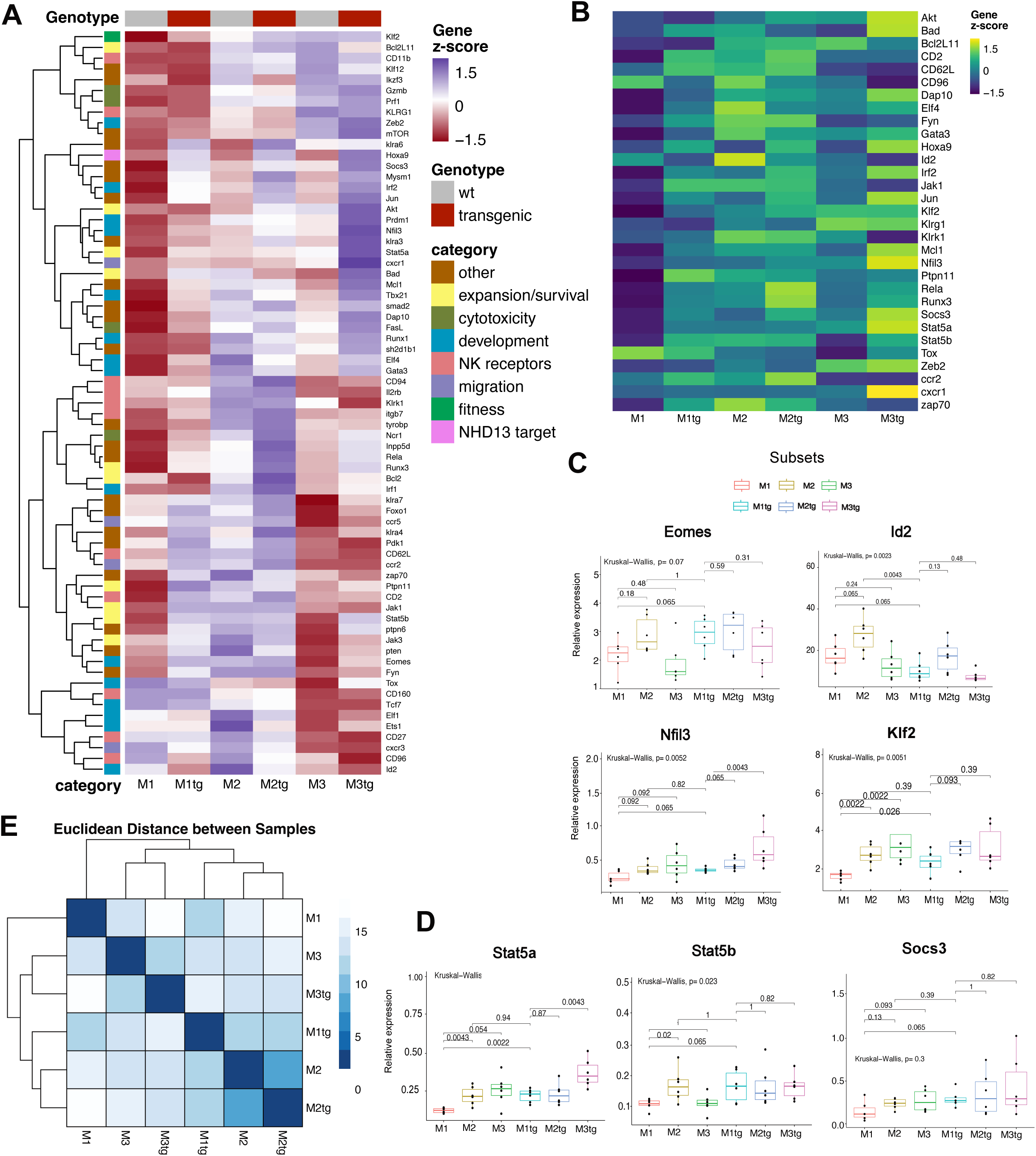
Gene expression profile of NHD13^tg^ NK cell subsets. **(A)** Heatmap of HPRT-normalized gene set from Fluidigm multiplexed RT-qPCR generated with the libraries ggplot2 (geom_tile) and ggdendro in R with low expressing gene in red colour-range, high expressing gene in blue colour-range and intermediate expression level in white. **(B)** Heatmap derived from Fluidigm main variable features. **(C-D)** Relative expression of genes involved in NK cell development **(C)** and NK cell signaling **(D)**. **(E)** Euclidean distance between samples. Mean comparison between subsets have been calculated with rstatix library on R. Each NK cell-subset gene analysis was a combination of 6 experimental replicates (3 replicates from a pool of 2 males per genotype and 3 replicates from a pool of 2 females per genotypes).

Altogether, these data suggest that M1 NK cells from NHD13^tg^ mice prematurely entered subsequent maturation/differentiation stage at the transcriptional level, and resemble the M2 stage of normal NK cell development.

### Potential mechanism behind MDS-induced NK cell deficiency and targets identification in NHD13^tg^ mice

The gene expression profiling highlighted the alterations in expression of key transcription factors Nfil3, Klf2, Id2 and Socs3 that control NK cell development, homeostatic proliferation and responsiveness to IL-15 (Figure 7A-D). To unravel potential unknown interaction among these transcription factors, we explored the ATAC-sequencing data-set generated by Yoshida *et al*.(*25*) and discovered that Nfil3 could likely induce Klf2 expression since its binding motif was found in close proximity to Klf2 (data not shown).

Based on this result, we established a probable network of regulatory pathways driven by *NUP98-HOXD13* fusion gene through the induction of Stat5 upregulation. The increased level of Stat5a detected in NK cells from NHD13^tg^ mice expression (Figure 7D, Supplementary Figure 9) leading to Nfil3 upregulation and the subsequent Klf2 upregulation. Since Id2 negatively regulates Socs3 expression, Id2 downregulation could result in the increased Socs3 expression (Figure 7C-D). These transcriptional changes together could contribute to the disease-induced NK cell defects observed in NHD13^tg^ mice (Supplementary Figure 9).

Finally, we extended the investigation of these gene candidates identified in NK cells from NHD13^tg^ mice into human settings, and measured the average gene expression in CD56^+^ mononuclear BM cells (BM-MNC) from healthy BM, MDS patients and AML patients with MDS-related mutations using a single cell RNA-seq data set (Figure 8). Interestingly, in a preliminary analysis, we found that *NFIL3* expression was higher, and *ID2* levels were lower, both, in MDS and AML patients with MDS-related mutations compared to the healthy donors, while *KLF2* expression was reduced in AML patients with MDS-related mutations, but not in MDS group (Figure 8). Although Socs3 level was altered in NHD13^tg^ mice, it did not show a clear diagnostic relevance in a human settings (Figure 8).

**Figure 8.**
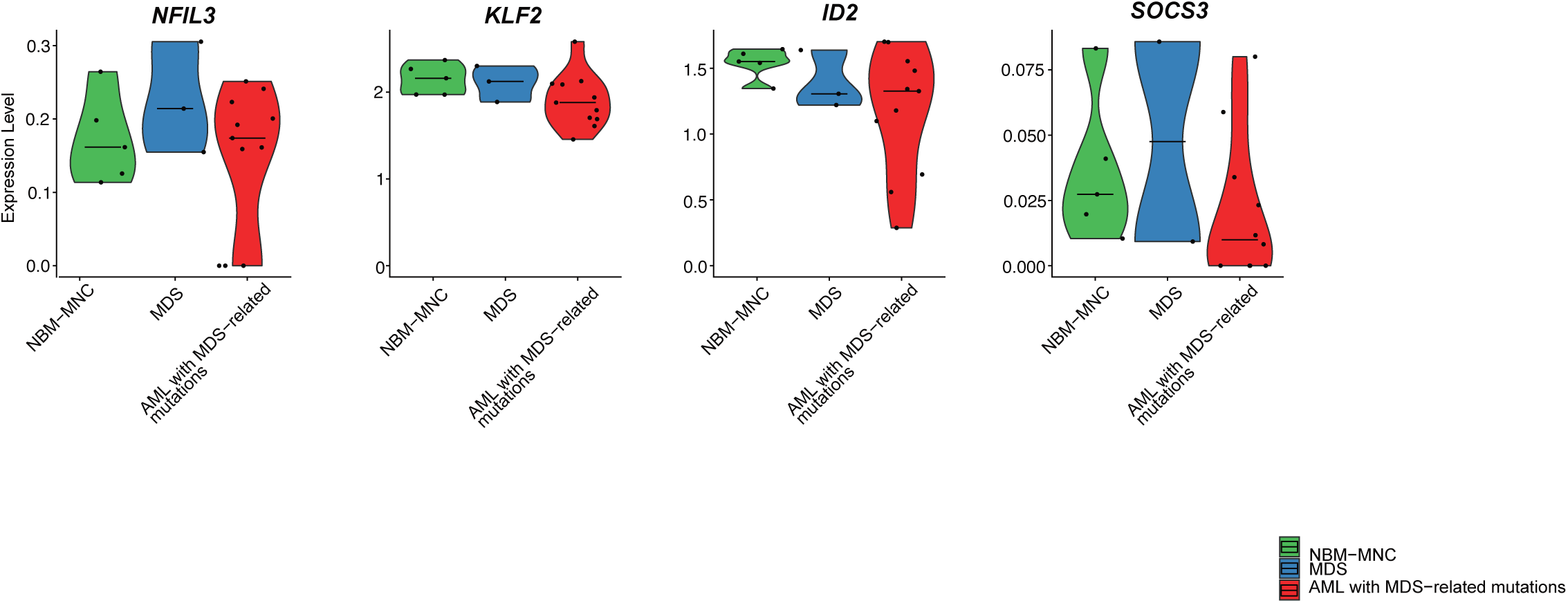
Expression of key NK cell regulatory genes is altered in patients with MDS and AML. Violin plots show the expression level of gene candidate *NFIL3*, *KLF2*, *ID2* and *SOCS3* in CD56^+^ bone marrow mononuclear cells (BM-MNC) from healthy individuals, MDS or AML patients with MDS-related mutations.

Taken together, we propose a regulatory network involving Nfil3, Klf2 and Id2 associated with the early reduction of the mNK cell pool at the pre-MDS stage in NHD13^tg^ mice that could be relevant as potential diagnostic genetic markers for MDS.

## DISCUSSION

Transgenic NHD13 mice expressing human *NUP98-HOXD13* fusion gene that is found in patients with myeloid malignancies, provide a valuable pre-clinical model to study the slow progressing disease from MDS to leukemia including AML and T-ALL (*12–14*) (*11*).

One of the limitations in modeling human disease in animals, is a transmission of somatic mutation into specific lineages. In MDS patients with monosomy 7, approximately 60% of NK cells carried chromosomal aberration (*26*), and in nearly all NK cells from patients with chronic myelogenous leukemia (CML) *BCR-ABL* fusion gene was detected (*27*). These findings together support that NHD13^tg^ model, where NK cells expressed the *NUP98-HOXD13* transgene, resembles the human disease.

The impact of leukemic transformation on the adaptive immune system was previously addressed in the NHD13^tg^ model, demonstrating the reductions in B and T cells (*12–14*). Importantly, selective T and B cell depletion after crossing NHD13^tg^ mice with RAG-1 knockout mice resulted only in a modest increase in leukemia frequency and decrease in the survival compared to NHD13^tg^ mice, suggesting a mild anti-leukemia effect of T cells and involvement of other immune cells (*15*). Since NK cells represent another lymphocyte population responsible for controlling cancer, which is maintained in RAG-1 knock-out setting, we here investigated how the disease onset and progression affects NK cell development, fitness and function, and whether specific NK cell loss impacts the disease progression.

Strikingly, selective NK cell depletion accelerated the progression from MDS to AML or T- ALL, leading to a decrease of overall survival of NHD13^tg^ mice treated with anti-NK1.1 antibody, the appearance of blasts up to 30% in the PB and severe lymphopenia in some cases, characteristic for leukemic progression. These results are the first direct evidence demonstrating the critical NK cell role in controlling MDS progression.

Interestingly, and not previously reported, we found that NHD13^tg^ mice had a significantly reduced pool of conventional NK cells in the PB and spleen clearly manifested already at the pre-MDS blast-free stage (6-9 week-old) before overt disease onset, progressing to a severe NK cell loss at the MDS stage (5-7 month-old). Approximately 20 % of NHD13^tg^ mice at the MDS-stage had no detectable NK cell levels, and similar as observed in patients with myeloid diseases (*7, 8*), a greater NK cell loss correlated with more advanced disease phenotype.

Sequential NK cell differentiation in the spleen of NHD13^tg^ mice was perturbed, leading to an accumulation of M1, a reduction in M2 mNK cells and a specific loss of mNK cells expressing KLRG1 receptor also detectable in the BM. Since KLRG1 receptor expression reflects the final NK cell maturation stage (*18*); the loss of KLRG1 positive NK cells in NHD13^tg^ mice further supports a developmental defect. Defects in NK cell maturation have been reported in AML patients, however the potential mechanisms responsible for this NK cell “hypomaturation” have not been established (*28, 29*).

The developmental alteration in NK cell compartments in NHD13^tg^ mice were cell autonomous, as BM from NHD13^tg^ mice specifically reconstituted the phenotype of donor NHD13^tg^ mice when competitively transplanted into WT recipients with no evident effect on co-transplanted WT BM competitor cells.

Although, the expression level of IL-2/IL-15 receptor β-chain (CD122) was not changed in NK cells from NHD13^tg^ mice, their response to IL-15 and IL-2 signaling was impaired as underlined by their poor ability to survive *in vitro*. In line with IL-15-response defect, the pool of KLRG1 positive NK cells, generated from M1 cells after homeostatic proliferation in response to IL-15 (*18*), decreased in NHD13^tg^ mice.

Perturbation in NK cell maturation in NHD13^tg^ mice can be, in part, due to the reduced level of Id2, that controls several aspects of NK cell development and IL-15 responsiveness (*23, 24*).

Decreased expression of Id2 transcripts and subsequent increase of Socs3 level in NK cell subsets from NHD13^tg^ mice could explain the impaired response to IL-15 through negative feedback induced by SOCS3-dempened IL-15 signal transduction. This possibility is supported by the partial rescue observed in the high IL-15 concentration in the competitive survival assay. Fauriat et al, recently suggested that a chronic cytokine stimulation involving the IL-15/mTOR pathway was responsible for the NK cell exhaustion leading to the reduction of NK cells and the loss of *in vitro* response to IL-15 signaling observed in leukemic mouse model (*30*).

Using gene profiling of NK cells at M1 and M2 developmental stages, we identified changes in the expression of several genes critical for NK cell development in NHD13^tg^ mice. Most of the transcription factors known to regulate NK cell maturation (Nfil3, Tox, Tcf7, Gata3, Irf2, and T-bet) (*21, 22, 31, 32*), and homeostatic proliferation (Klf2) (*33*) were expressed at higher levels in M1 NK cells from NHD13^tg^ mice that were compatible with the M2 maturation stage. Although M1 NK cells prematurely entered the next developmental stage at the transcriptional level, this was not accompanied by the increase of functional maturation expected to be acquired by M2 NK cells from NHD13^tg^ or WT littermate mice during development.

The reduced *ID2* expression, both in patients with myelodysplastic malignancies and AML patients with mutations related to MDS, and decreased *KLF2* levels in AML patients, could potentially explain NK cell exhaustion through maturation and proliferation defects. This highlights their potential application as new genetic markers in the diagnosis of early MDS disease stage. Studies by Vivier et al (*34*), reported an increased *KLF2* levels using a single cell RNA sequencing of NK cells from leukemia patients. This difference in *KLF2* expression pattern could be explained by the fact that AML cases analyzed in that study were normo-cellular for the NK cell subset, while our work focused on disease with a clear NK cell reduction. It has been previously reported that MDS and AML patients can either be normo-cellular or hypo-cellular with an “hypo-mature” profile (*35*). Importantly, previous work pointed-out the prognostic value of the size and the function of NK cell population in AML patients in predicting their response to the therapy (*36–38*). Combining measurement of NK cell pool with monitoring the expression level of *ID2* and *KLF2* in isolated NK cells could provide an indication of the disease progression.

## MATERIALS AND METHODS

### Mice

Vav1-*NUP98/HOXD13* transgenic (NHD13^tg^) (*12*) and B6SJL CD45.1 mice on C57Bl/6 background were on obtained from The Jackson Laboratory.

### Tissues

Bone marrow cells (BM) were extracted from tibias and femoras using mortar and filtered using a 100-µm nylon mesh filter. Spleens were isolated and smashed through a 100-µm mesh using a syringe piston, then washed and splenocytes were filtered using a 70-µm nylon mesh filter (BD). Peripheral blood cells (PB) were collected from tail vein into heparin tubes and ACK (Ammonium-Chloride-Potassium) lysis buffer treatment was used to remove erythrocytes. Single cell suspensions from all extracted tissues were prepared in PBS 2 mM EDTA 5 % FCS and cells were counted using Sysmex (KX-21N) Hematology analyzer.

### Antibodies, Flow cytometry and Cell sorting

Cells were stained for 40 min in PBS supplemented with 2mM EDTA and 5 %FCS, on ice protected from light with specific mAb (**Supplementary Table I**). Blood cell distribution was evaluated using anti-mouse antibodies targeting specific lineage surface markers: CD3^+^NK1.1^-^ (T cells), CD19^+^ (B cells), NK1.1^+^CD3^-^ (NK cells), CD11b^+^Gr1^+^ (myeloid cells) and cKIT^+^ or CD11b^+^cKIT^+^ (blast cells), and Ter119^+^ (erythroid cells). Peripheral conventional NK cell subsets were identified using Chiossone et al. (*17*) phenotypic definition: Lineage negative (Lin) (CD19, CD11b, Gr1, Ter119)^-^CD3^-^CD122^+^NK1.1^+^DX5^+^CD27^+^CD11b^-^ (M1), Lin^-^CD3^-^ CD122^+^NK1.1^+^DX5^+^CD27^+^CD11b^+^ (M2) and Lin^-^CD3^-^CD122^+^NK1.1^+^DX5+CD27^-^CD11b^+^ (M3) cells. BM lymphoid progenitors were defined according to Fathman et al. (*16*) phenotypic definition: common lymphoid progenitor (CLP) as Lin^-^ CD27^+^CD244^+^CD127^+^cKIT^+^CD135^+^CD122^-^, (pre-NKP) as Lin^-^ CD27^+^CD244^+^CD127^+^cKIT^+^CD135^-^CD122^+^, and NKP as Lin^-^ CD27^+^CD244^+^CD127^+^cKIT^+^CD135^-^CD122^+^ cells. To exclude dead cells, 7- aminoactinomycin D (7-AAD) (Sigma-Aldrich) was used. Intracellular staining of IFN-γ was performed using Cytofix/Cytoperm permeabilization kit (BD). Fluorescence minus one (FMOs) were used to determined positive signal and compensation control together with single-stained mAbs. All samples were acquired on Fortessa (BD) and analyzed with FlowJo analysis software (TreeStar). Cell sorting was performed on FACS AriaIIu or AriaIII (BD) using the purity precision mode.

### In-vivo NK cell depletion

A cohort of 6 NHD13^tg^ mice at 10 to 11 months old (late MDS stage with a risk of progressing to leukemia) were treated with a single intra-peritoneal injection of anti-NK1.1 antibody (200 µg) (NordicBioSite) to induce the depletion of mature NK cells (*19*).

PB lineage distribution was evaluated before treatment, 1 months and 3 months after treatment. A cohort of 3 age-matched NHD13^tg^ mice were treated with anti-IgG2aκ antibody (200 µg) (NordicBioSite) as a negative control group. Non-treated WT littermate mice were used as a negative reference and an age-matched non-treated NHD13^tg^ mouse at the leukemia stage was used as a positive reference.

### Functional NK cell analysis /NK cell degranulation assay and IFN-γ production

NK cell cytotoxic killing and IFN-γ production were evaluated by plating 2.5 x 105 unfractionated splenocytes from NHD13^tg^ or WT littermate mice in 100 µL per well into U-bottom 96-well plates. NK cells were specifically activated with purified anti-mouse NK1.1 mAb (7 µg/mL) and/or a combination of murine recombinant mIL-12 (10 ng/mL) and mIL-18 (50 ng/mL) together with Monensin (1X) and Brefeldin A (1X) (BioLegend) to detect cytotoxicity and IFNγ production, respectively. Cells activated with PMA/Ionomycin (PMA/ION) were used as positive control and untreated cells as a negative control. Ten wells were pooled for the analysis of each condition. Cells were harvested after 5h incubation at 37°C and stained for detection of cell surface expression of CD107a or intracellular IFN-γ expression within the NK cell compartment using flow cytometry.

### NK cell response to IL-2 and IL-15

Splenocytes from NHD13^tg^ and WT littermate controls were labeled with two intracellular dyes: Cell Trace Far Red (CTR) and Cell Trace Violet (CTV) (Invitrogen) respectively, plated at a ratio of 1:1, and cultured for 7 days in the presence of cytokines. Expression of CTR or CTV in remaining NK cells was assessed by flow cytometry.

### Competitive transplantation assay

Chimera mice were generated by transplanting a 1:1 mixture of unfractionated BM cells from CD45.2 NHD13^tg^ mice or CD45.2 wild type (WT) littermates together with CD45.1 WT B6SJL BM cells into lethally irradiated (900 cGy) CD45.1 WT B6SJL recipient mice. A total number of 1×10^6^ cells was transplanted per mice via intravenous injection and PB cells were analyzed at 4-, 6- and 8-months post-transplantation to assess donor-derived multi-lineage reconstitution. Splenic blast level was evaluated in all remaining recipient mice euthanized at 8-months post-transplantation.

### Gene expression analysis by RT-qPCR

Cells were sorted in triplicates, at 20 cells per well (96-well plates) into 4 μL of lysis buffer containing 0.4% NP40, deoxynucleoside triphosphates, dithiothreitol, and RNase OUT (Invitrogen), and were snap frozen. RT-PCR and 18 cycles of pre-amplification with a mix of Taqman probes (at 0.4x) was performed in each well following the One-Step qRT/PCR with ROX protocol (Invitrogen). The diluted product and Taqman probes (**Supplementary Table II**) were loaded into a 96.96 Dynamic Array IFC (Fluidigm) following the manufacturer’s protocol. Chips were run in Biomark (Fluidigm) and data analyzed through RT-PCR Analysis Software (Fluidigm). B2m and Hprt were used as reference genes for normalization. Relative expression Log10 (2-ΔCt) of each gene was used to create a normalized heatmap using the command geom_tile and ggdendrogram from ggplot2 and ggdendro libraries in R software.

### Re-analysis of M1 and M3 NK-cell ATAC-sequencing data

ATAC-sequencing raw data from BM M1 (CD27^+^CD11b^-^) and M3 (CD27^-^CD11b^+^) mature NK cell were downloaded from the GEO database (GSE100738) (*25*). Briefly, paired-end reads were trimmed using Trimmomatic and mapped to mm10 (Gencode build GRCm38.p6) with Bowtie2 (v. 2.3.4.2) (*39*). Mitochondrial reads were filtered out with Samtools (v. 1.9) followed by PCR duplicate and low-quality sequence (MAPQ < 10) removal with (Picard MarkDuplicates (v. 2.15.0)) and Samtools, respecively. Peak calling from the mapq10 filtered bam file was performed using the Kundaje lab ATAC-seq pipeline (https://www.encodeproject.org/pipelines/ENCPL792NWO/) with MACS2 and IDR (irreproducible discovery rate) threshhold set to 0.05 and 0.1, respectively. From each processed duplicate, the Optimal peak set was used for downstream analysis. Additionally, the mapq10 filtered bam files used as input for peak calling were also used to create Tag directories with the HOMER platform (v. 4.10) (*40*). Peak files from the individual samples were combined with Bedtools intersectBed (*41*). For analysis of dynamic chromatin changes, read counts were derived in peak regions using Homer (annotatePeaks.pl –noadj) and tested for differential peak accessibility with edgeR (v. 3.26.8) (*42*). Bioconductor package via HOMER getDifferentialExpression.pl. Differential peaks with adjusted p-value ≤ 0.05 and |F.C| ≥ 2 were considered significantly different.

To identify potential regulators of the Klf2 gene, we used HOMER annotatePeaks.pl to assign Klf2 coordinates to the mm10 genome and then extended these coordinates with 2Mb in both directions. The extended region was then overlapped with M1/M3 differential ATAC-seq peaks using Bedtools intersectBed and screened for motifs from the JASPAR2022 database using gimme scan (with an 1 % FPR) from the GimmeMotif (v. 0.17.0) software suite (*43*) to generate a log odds score for each motif in each accessible region.

### Single-cell RNA-sequencing

Mononuclear cells from three MDS samples, eleven AML samples harboring MDS-related mutations, and five human normal bone marrow samples were characterized using 10X genomics chromium 3’ v3 single cell sequencing as part of a larger AML study (Lilljebjörn et al, unpublished). In brief, single cell sequencing libraries were prepared and sequenced in accordance with the manufactureŕs instructions. The single cell data were preprocessed using Cellranger (v. 3.1.0; 10X genomics) and analyzed using Seurat (v. 4.0.0) (*44*). Cell type prediction was performed using Seurats TransferData function based on a human bone marrow reference dataset (*45*). The average gene expression level for each cell type in each sample was calculated using Seurats AverageExpression function. From this data, the average expression level for NFIL3, KLF2, ID2 and SOCS3 was visualized using ggplot2.

### Study approval

The Ethical Committee at Lund University approved all animal studies performed.

The collection of human material was done after written informed consent according to the Declaration of Helsinki with approval of the Local Ethical Committee.

### Statistics

Graphs and statistical analysis were performed using GraphPad 8 (GraphPad Software, La Jolla, CA). The statistical significances between groups were determined by the Mann-Whitney U test for unpaired samples and ANOVA multiway comparison.

## ACKNOWLEDGMENTS

We thank Anna Fossum, Zhi Ma and Mickael Sommarin at Lund Stem Cell Center FACS core facility for their expertise. We thank Mattias Carlsten for critical review of the manuscript. This work was supported by grants from the Swedish Cancer Society, the Swedish Research Council and the Alfred Österlunds Foundation (awarded to ES).

## Author contributions

ES conceived, designed and supervised the study with contributions from DB, CB and TF. GT- D designed and performed the experiments with contributions from OY, OK, MC, JU, DNV and HL. GT-D, ES and HL analyzed and interpreted data. GT-D and ES prepared the figures and wrote manuscript with input from all authors.

