## Supplementary Information for "NK cells control the progression of myelodysplastic syndrome but become initial disease target in *NUP98-HOXD13* mouse model"

**Supplemantary Figures 1-9**

**Supplementary Table I and Table II**

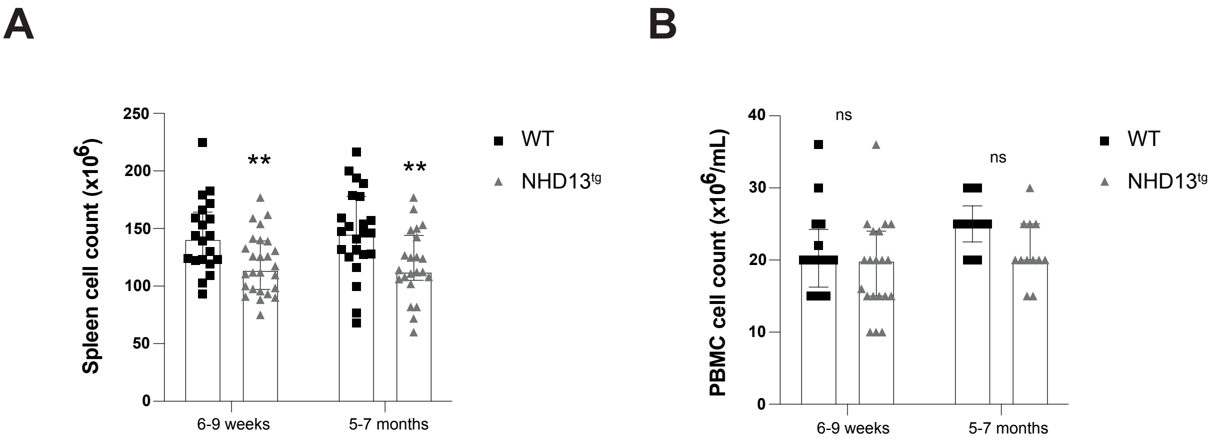


**Supplementary Figure 1 (related to Figure 2). Tissue cellularity in NHD13$^{tg}$ mice during MDS progression.** Total number of **(A)** splenic and **(B)** peripheral blood mononuclear (PBMC) cells. Each bar represents the median with value with the interquartile range from 6-9 weeks and 5-7 months old NHD13$^{tg}$ mice (grey triangles) and WT littermate controls (black squares). Ten to twenty mice were analyzed per group. Statistical differences between the groups were determined using unpaired Mann-Whitney U test

**p ≤0.01, ns= not significant.

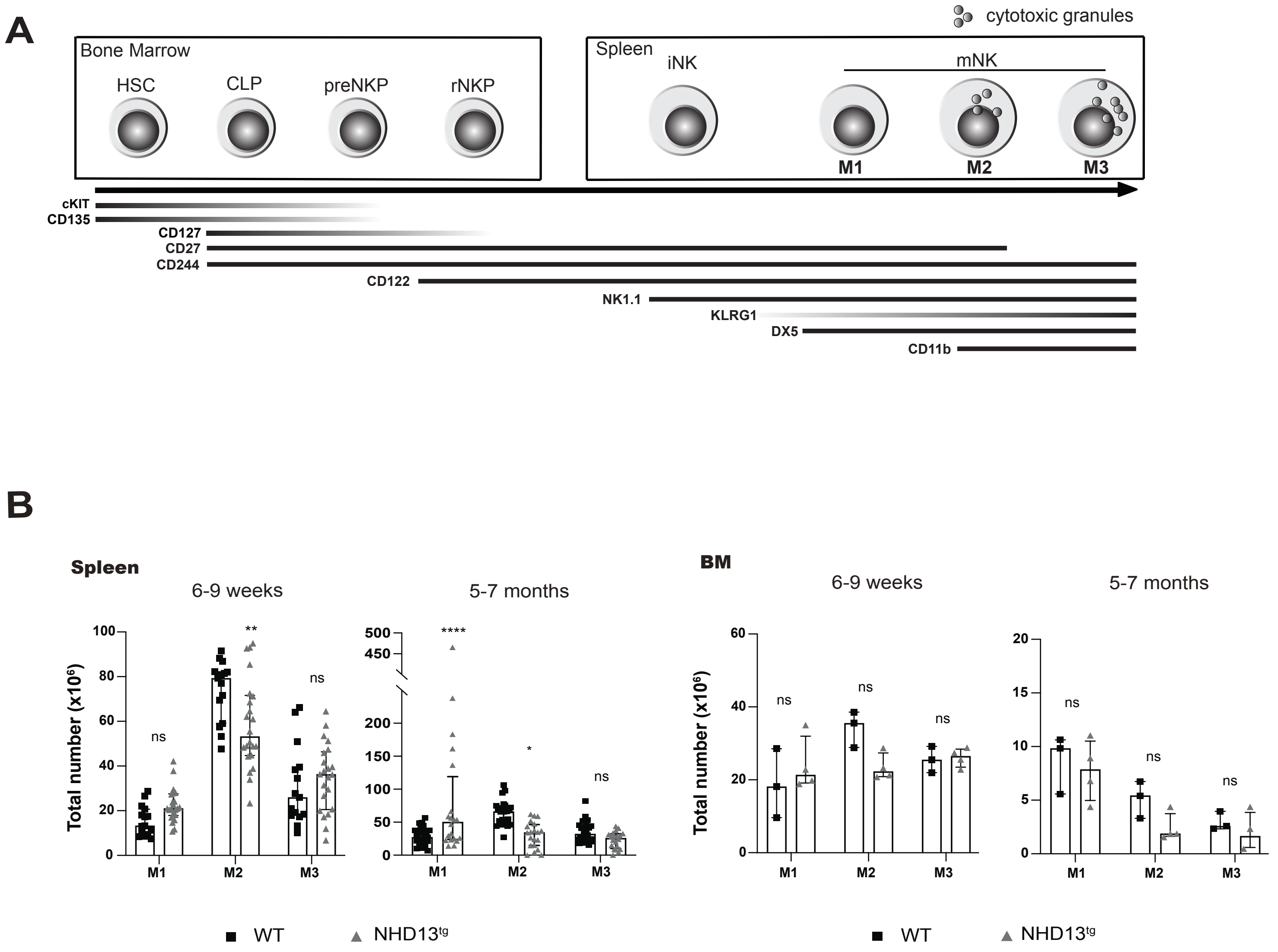


**Supplementary Figure 2 (related toFigure 3). Perturbed NK cell development in ND13$^{tg}$ mice.**

**(A)** Schematic overview of mouse conventional NK cell development from hematopoietic stem cells (HSCs) in the bone marrow (BM), and the surface markers defining each sequential stage including: CLP (Lin$^{-}$CD27$^{+}$ CD244$^{+}$CD127$^{+}$cKIT$^{+}$CD135$^{+}$CD122$^{-}$), pre-NKP (Lin$^{-}$CD27$^{+}$CD244$^{+}$CD127$^{+}$cKIT$^{+}$CD135$^{-}$CD122$^{+}$), and NKP (Lin$^{-}$CD27$^{+}$CD244$^{+}$CD127$^{+}$cKIT$^{+}$CD135$^{-}$CD122$^{+}$). Conventional NK cell maturation continues in the spleen with iNK (CD122$^{+}$NK1.1$^{+}$DX5$^{-}$) and mature NK cells (mNK) (CD122$^{+}$NK1.1$^{+}$DX5$^{+}$) pool containing M1 (CD27$^{+}$ CD11b$^{-}$), M2 (CD27$^{+}$CD11b$^{+}$) and M3 (CD27$^{-}$CD11b$^{+}$) subsets. Terminally differentiated M2 and M3 subsets are functionally mature as they contain cytotoxic granules. **(B)** Total number of mature NK cells in the spleen (left panels) and BM (right panels) in 6-9 weeks and 5-7 months old NHD13$^{tg}$ mice (grey triangles) and WT littermate controls (black squares). Each bar represent the median value with the interquartile range, 10-20 mice were analyzed in the spleen group and 3-4 mice in the BM group. Statistical differences were determined using Two-Way ANOVA multiple comparisons *$p \leq 0.05$, **$p \leq 0.01$, ****$p \leq 0.0001$, ns= not significant.

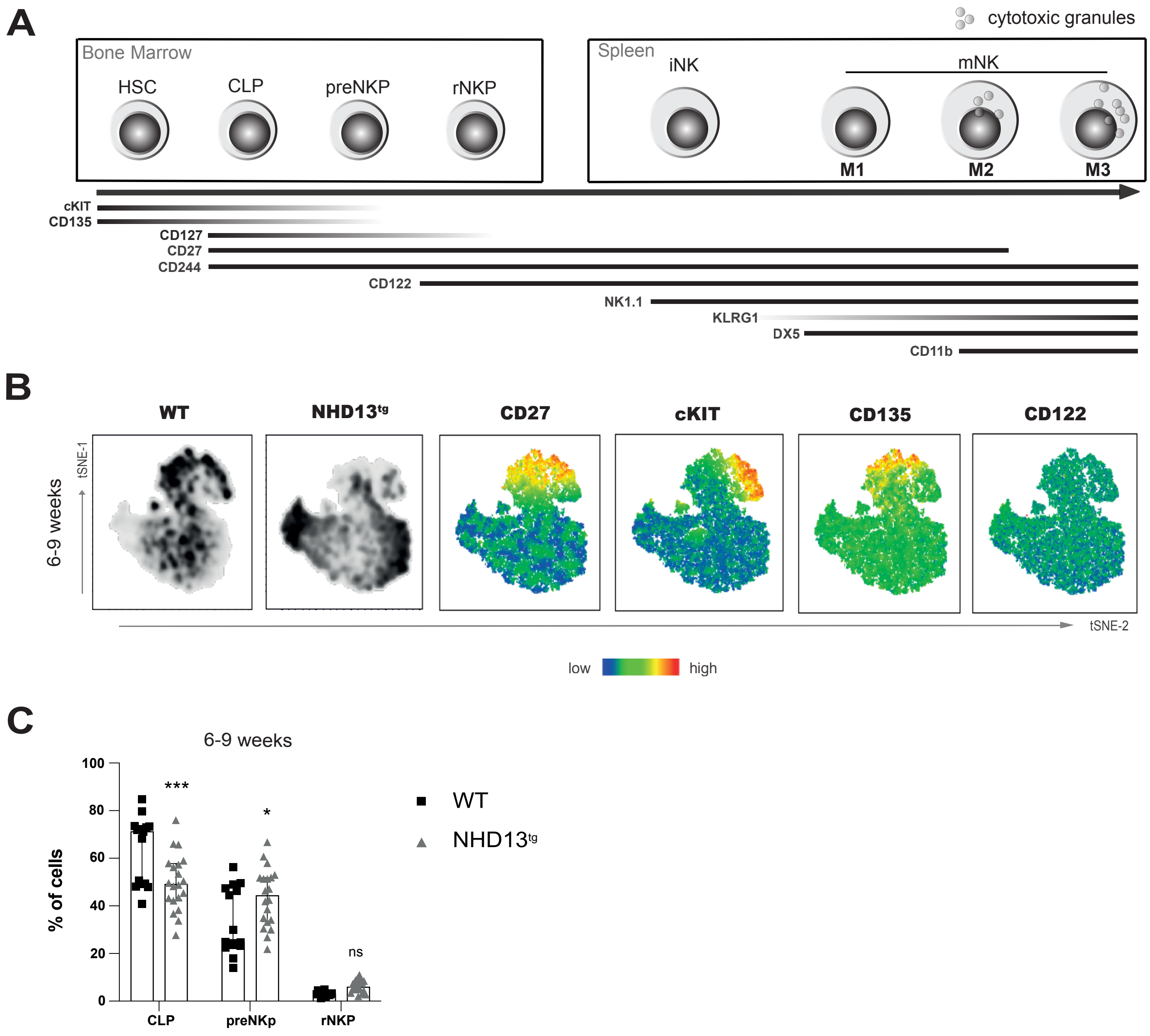


**Supplementary Figure 3 (related to Figure 3). Development of NK cell BM progenitors.**

**(A)** Schematic overview of mouse conventional NK cell development from hematopoietic stem cells (HSCs) in the bone marrow (BM), and the surface markers defining each sequential stage including: CLP (Lin$^-$CD27$^+$ CD244$^+$CD127$^+$cKIT$^+$CD135$^+$CD122$^-$), pre-NKP (Lin$^-$CD27$^+$CD244$^+$CD127$^+$cKIT$^+$CD135$^-$CD122$^{+}$), and NKP (Lin$^-$CD27$^+$CD244$^+$CD127$^+$cKIT$^+$CD135$^-$CD122$^+$). Conventional NK cell maturation continues in the spleen with iNK (CD122$^+$NK1.1$^+$DX5$^-$) and mature NK cells (mNK) (CD122$^+$NK1.1$^+$DX5$^+$) pool containing M1 (CD27$^+$CD11b$^-$), M2 (CD27$^+$CD11b$^+$) and M3 (CD27$^-$CD11b$^+$) subsets. Terminally differentiated M2 and M3 subsets are functionally mature as they contain cytotoxic granules.

**(B)** Representative t-distributed stochastic neighbour embedding (t-SNE) density map of BM samples from 6-9 weeks and 5-7 months old NHD13$^{tg}$ mice and WT littermate controls. Heatmap statistics shows the expression level of keys surface markers CD27, c-KIT, CD135 and CD122 allowing the localization of early progenitors within the map. **(C)** Percentage of different BM progenitors in 6-9 weeks and 5-7 months old NHD13$^{tg}$ mice (grey triangles) and WT littermate controls (black squares). Each bar represents the median with value with the interquartile range. Ten to twenty mice were analyzed per group. Statistical differences between the groups were determined using Two-Way ANOVA multiple comparisons *p ≤0.05,***p ≤0.001, ns= not significant.

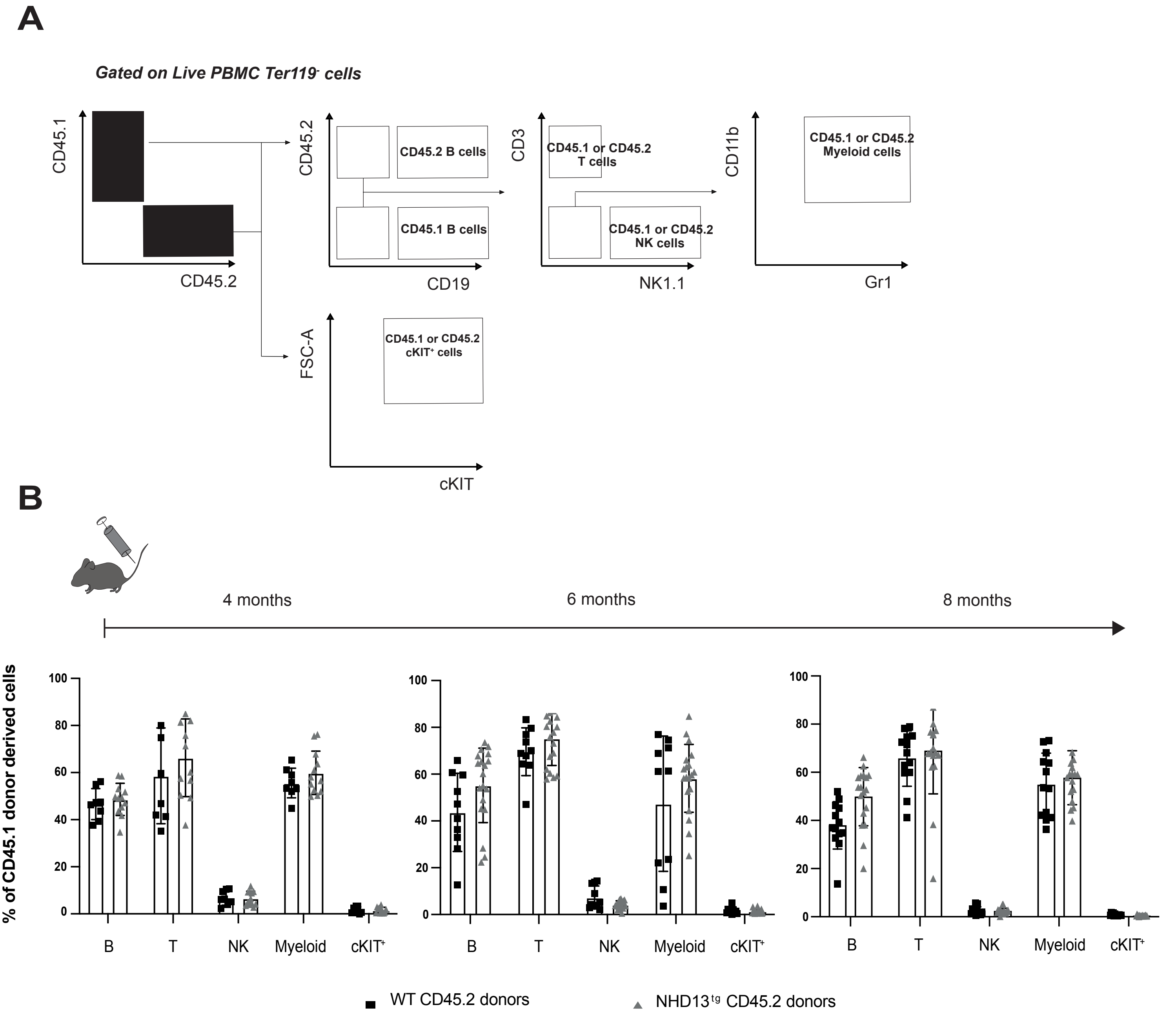


**Supplementary Figure 4 (related to Figure 4). CD45.1 donor-derived lineage reconstitution in chimera mice after competitive transplantation. (A)** Schematic overview of the gating strategy used to identify blood lineages derived from CD45.1 or CD45.2 bone marrow donor cells. Expression of different CD45 alleles was used to distinguish donor cells (CD45.2) from competitor or recipient cells (CD45.1). For each CD45 expressing cell type, blast cells ($cKIT^{+}$) and B ($CD19^{+}$) cells were first gated out. Next, the gate was set to identify T ($CD19^{-}CD3^{+}NK1.1^{-}$) and NK ($CD19^{-}CD3^{-}NK1.1^{+}$) cells. The final gating defined myeloid ($CD3^{-}NK1.1^{-}CD11b^{+}Gr1^{+}$) cells. (**B**) Percentage of CD45.2 $NHD13^{tg}$ (grey triangles) or WT littermates (black squares) donor cells together with CD45.1 WT competitor cells from B6SJL mice at 4, 6 and 8 months post-transplantation. Data are from 3 independent competitive transplantation experiments. At least 8 recipient mice were transplanted per group. Statistical differences were determined using Two-Way ANOVA with multiple comparison, ns= not significant.

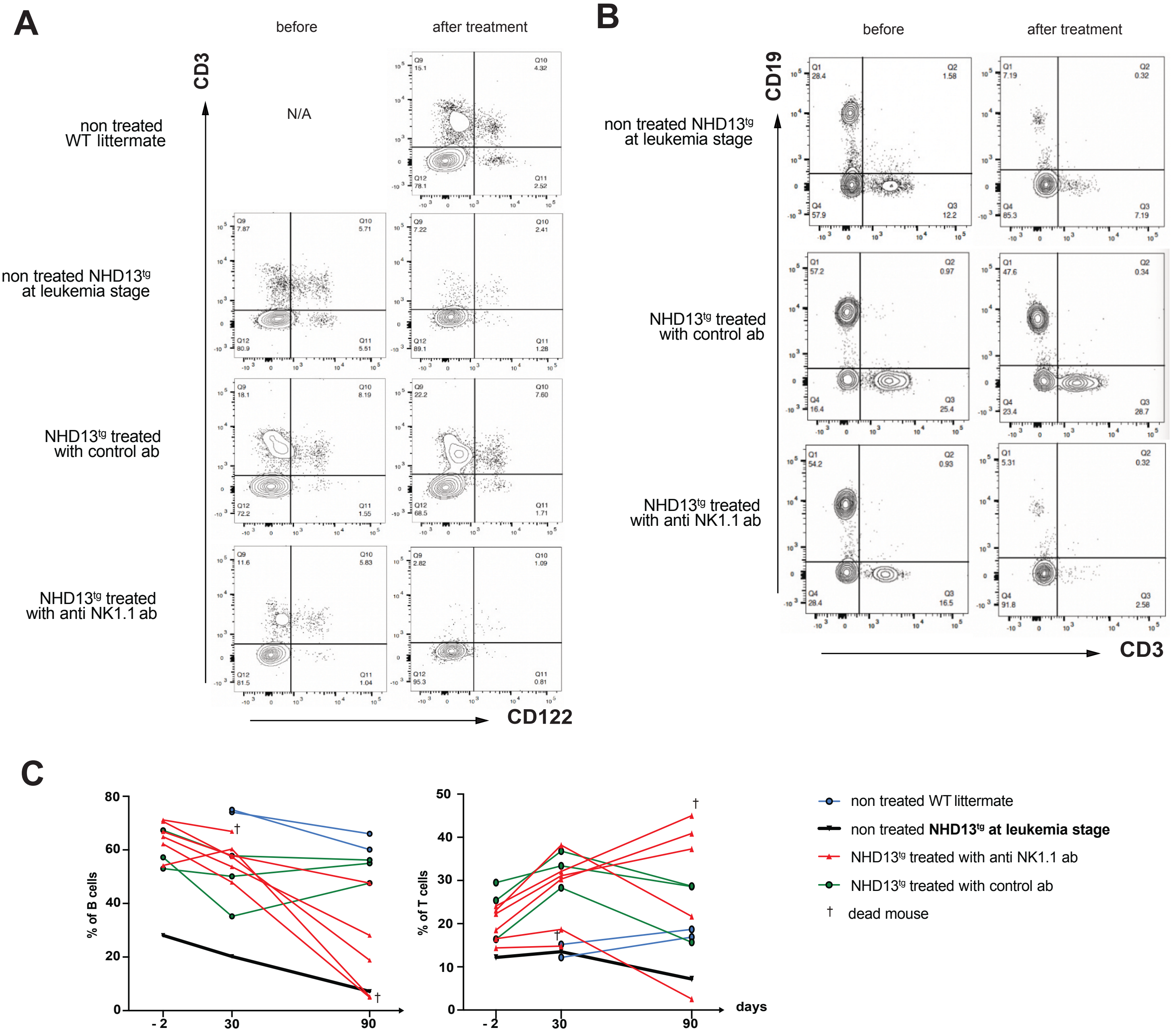


**Supplementary Figure 5 (related to Figure 5). Lymphoid compartment following selective NK cell depletion.**

**(A)** FACS™ profile of NK cell pool (CD3$^-$CD122$^+$) before and 3 months after treatment with anti-NK1.1 antibody or isotype control antibody. **(B)** FACS™ profile of lymphoid cells (CD19$^+$ or CD3$^+$) before and 3 months after treatment. **(C)** Percentage of B cells (CD19$^+$) and T cells (CD3$^+$) before (-2days) and after (30 and 90 days) specific NK cell depletion. A cohort of 10 to 11 months old NHD13$^{tg}$ mice injected with isotype control antibody was used as a negative control. Age-matched WT littermate mice were used as a negative reference, and a 1-year old NHD13$^{tg}$ mouse at the myeloid leukemia stage was used as a positive control. N/A= non applicable.

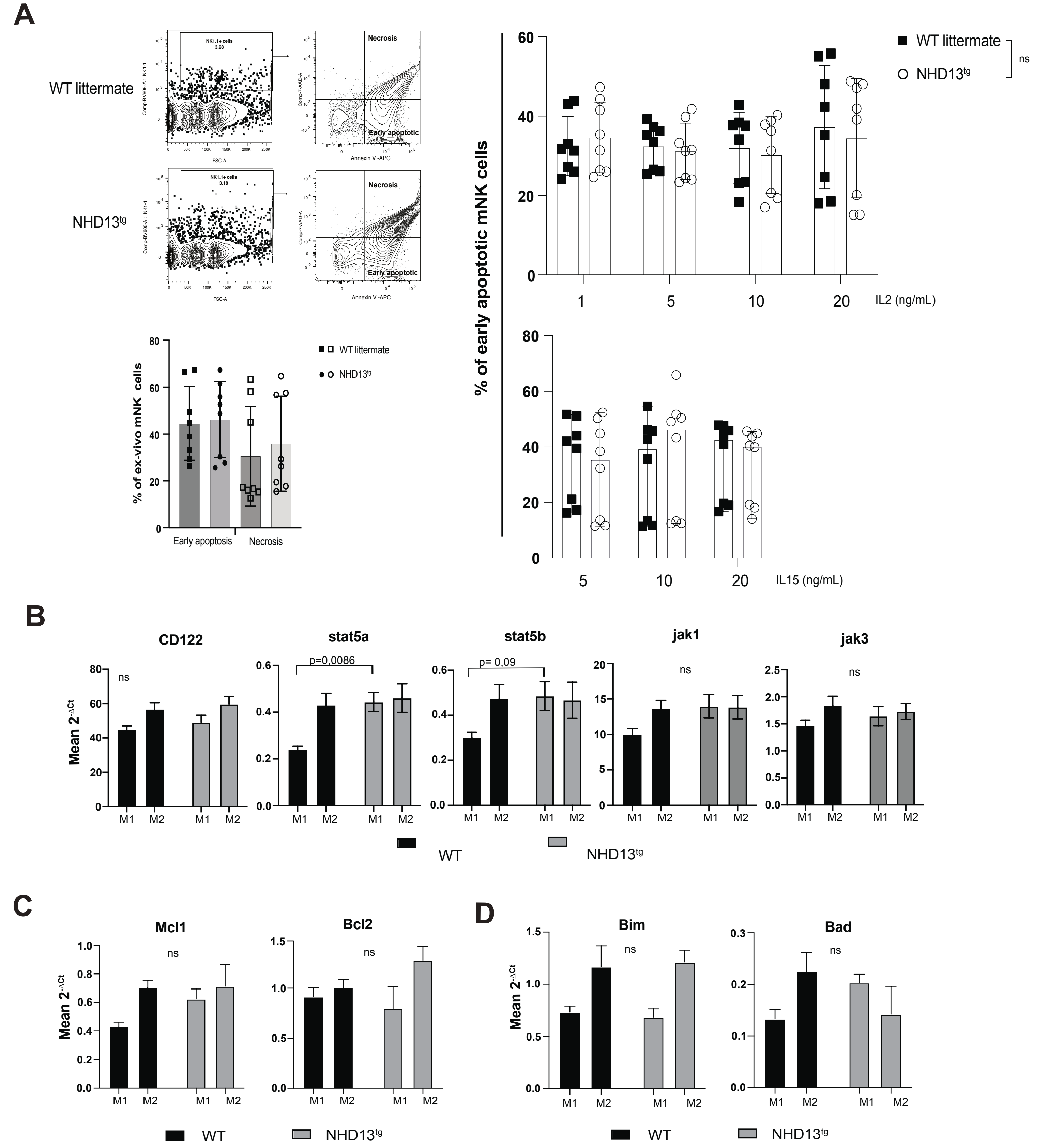


**Supplementary Figure 6. NK cell survival in response to IL-15 and IL-2 is not perturbed in NHD13$^{tg}$ mice.** **(A)** Representative FACS™ contour plot of early apoptotic (AnnexinV$^{+}$ 7AAD$^{-}$ NK1.1$^{+}$) mNK cells (top left panel) from spleen. Summary bar plot from ex-vivo analyzed splenocytes (bottom left panel) or after IL-2 (top right panel) and IL-15 (bottom right panel) stimulation. **(B-D)** Relative expression level of mRNA of **(B)** CD122 and transcripts encoding IL-2 and IL-15 downstream signaling targets, **(C)** anti-apoptotic, and **(D)** pro-apoptotic members of the Bcl2 family. All relative gene expressions were obtained from Fluidigm multiplex RT-qPCR analysis of sorted M1 and M2 mNK cell subsets from NHD13$^{tg}$ (grey bars) or WT littermate (black bars) mice. Bars represent the mean of the $2^{-\Delta Ct}$ replicate values with SEM normalized with each subsets Hprt1 respective Ct values. Nine mice were analyzed per group in 3 independent experiments. Statistical differences were determined using Two-Way ANOVA with multiple comparison * p≤0.01, ** p≤0.02, ns=not significant.

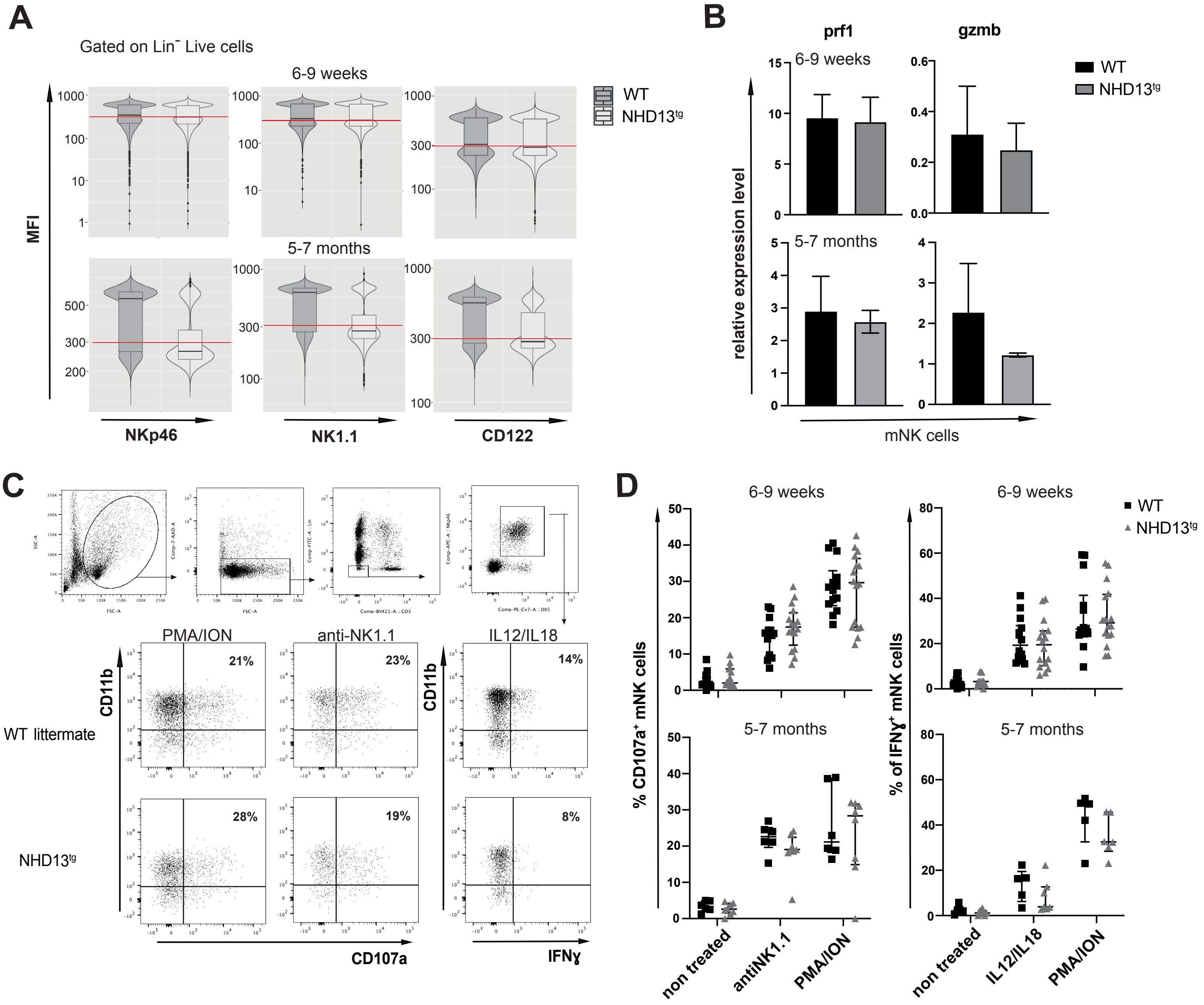


**Supplementary Figure 7. Mature NK cells in NHD13$^{tg}$ mice have normal cytotoxic function and cytokine production**. **(A)** Violin and box plots represent the median fluorescence intensity expression of NK cell receptors NKp46, NK1.1 and CD122 within Lin$^{-}$ Live splenocytes from 6-9 weeks and 5-7 months old NHD13$^{tg}$ (light grey) in comparison to WT littermate control (dark grey) mice. The red line represents the threshold of the positive signal based on each FMO parameter. **(B)** Relative expression level of Perforin (prf) (left) and Granzyme b (gzmb) (right) mRNA transcripts obtained from Fluidigm multiplex RT-qPCR analysis of sorted mNK cells from NHD13$^{tg}$ (grey bars) in comparison to WT littermate control (black bars) mice. Bars represent the mean of the $2^{-\Delta Ct}$ replicate values with SEM normalized with each subsets Hprt1 Ct values. **(C)** Gating strategy and representative FACS™ plots of splenocytes from 6-9 weeks old WT littermate (top) and NHD13$^{tg}$ mice (bottom) after stimulation with PMA/Ionomycine (PMA/ION) (left panel), anti-NK1.1 (middle panel) or IL-12/IL-18 (right panel). Cytotoxic response was measured with the upregulation of surface protein CD107a and cytokine production was evaluated by measuring the intra-cellular IFN-ɣ expression. **(D)** Percentage of mNK cells expressing CD107a (cytotoxic response) or IFN-ɣ (cytokine release) in response to different stimuli from 6-9weeks or 5-7 months old NHD13$^{tg}$ mice (grey triangles) and WT littermates (black squares). At least 6 mice were analyzed per group in 3 independent experiments.

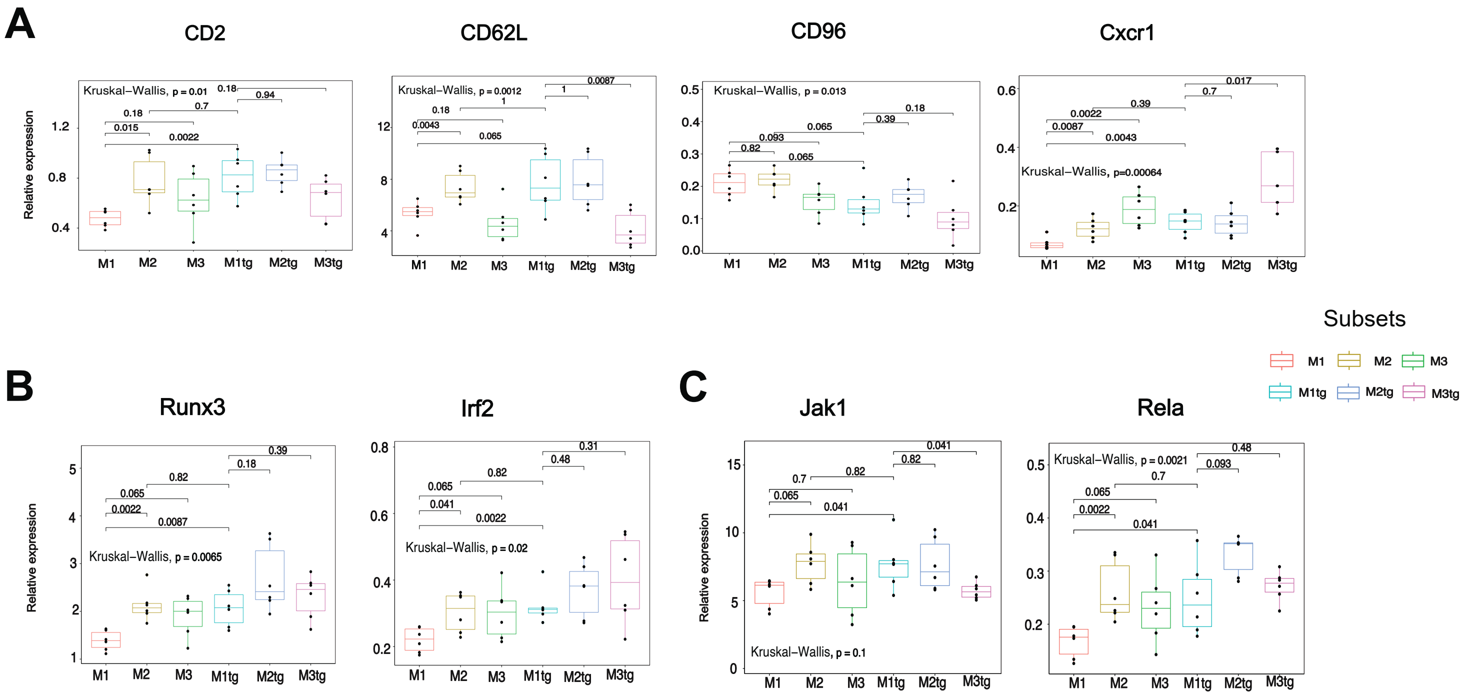


**Supplementary Figure 8 (related to Figure 7). Gene expression profile of NHD13$^{tg}$ NK cell subsets. (A-C)** Boxplots represent relative expression of genes **(A)** encoding cell surface markers, **(B)** genes regulating NK cell development, (**C**) genes involved in cell signaling. Mean comparison between subsets have been calculated with rstatix library on R.

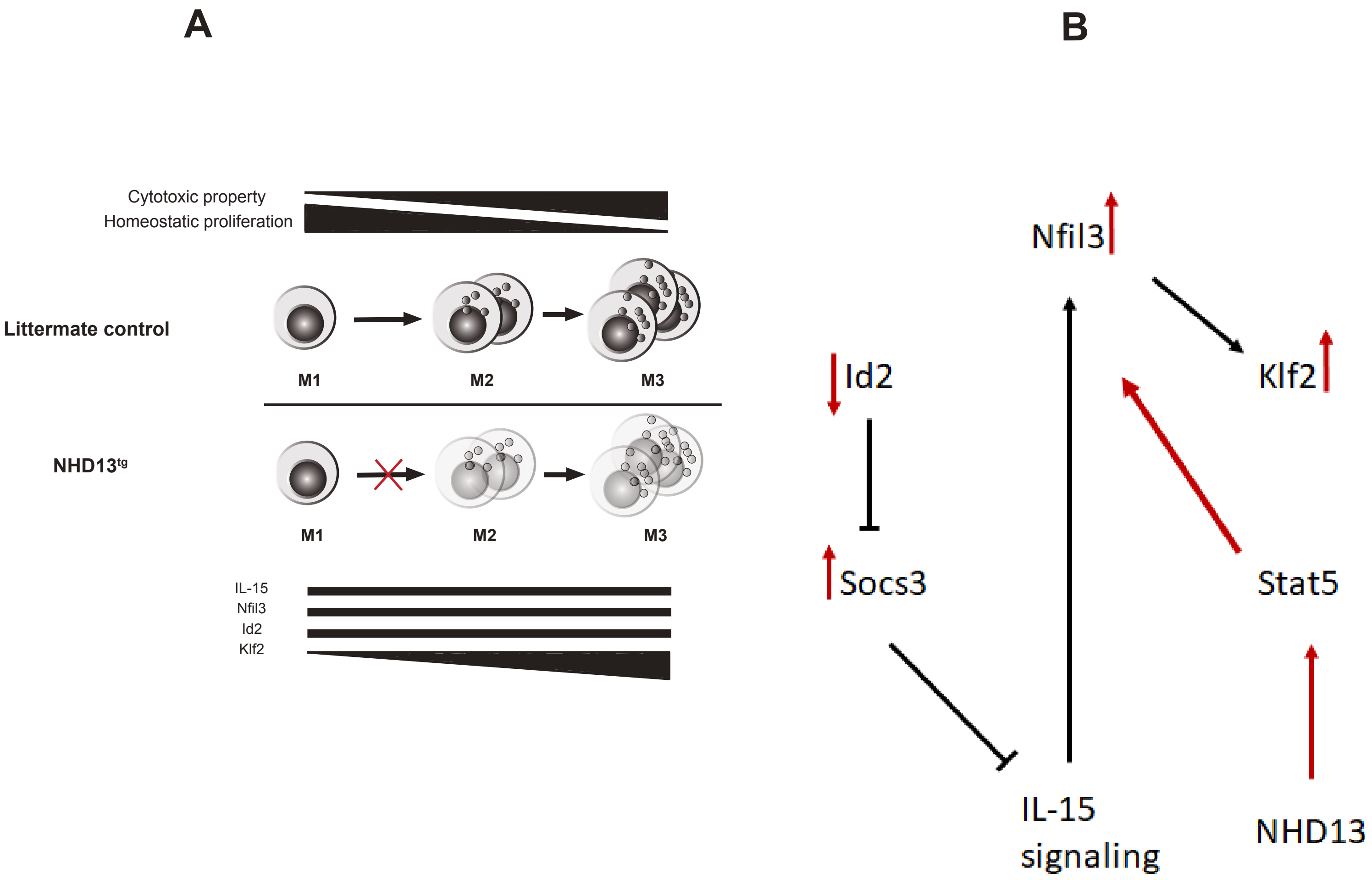


**Supplementary Figure 9 (related to Figure 7). Gene candidate and potential mechanism behind MDS-induced NK cell defects in NHD13$^{tg}$ mice.**

**(A)** Potential mechanism responsible for MDS-induced NK cell defects with a schematic overview of conventional mouse NK cell developmental stages in the periphery of littermate control mice and hypomaturation profile in NHD13$^{tg}$ mice. Terminally differentiated M2 and M3 subsets are functionally mature as they contain cytotoxic granules and have a decreased potential of homeostatic proliferation as the expression level of the transcription factor Klf2 increases. **(B)** Arrows show the changes in the expression level of Nfil3, Stat5a, Klf2, Id2 and Socs3 genes in NHD13$^{tg}$ mice compared to WT controls (see Figure 7) and indicate potential interactions within NK cell regulatory network.

SUPPLEMENTARY TABLES

### Supplementary Table I. List of Antibodies used for Flow Cytometry Analysis

| Antigen | Clone | Fluorochrome | Company |
|---|---|---|---|
| AnnexinV | - | AF647 | BioLegend |
| CD107a (LAMP-1) | 1D4B | PE | BD Biosciences |
| CD117 (c-kit) | 2B8 | APC/eFluor780 | eBioscience |
| CD117 (c-kit) | 2B8 | PE/Cy7 | BioLegend |
| CD11b (Mac-1) | M1/70 | FITC | eBioscience |
| CD11b (Mac-1) | M1/70 | APC/Cy7 | BD Biosciences |
| CD122 | TM-beta1 | PerCP/eFluor710 | eBioscience |
| CD127 | A7R34 | APC/Cy7 | BioLegend |
| CD19 | 6D5 | PE/Cy7 | BioLegend |
| CD19 | 6D5 | FITC | BioLegend |
| CD244.2 | 244F4 | APC | eBioscience |
| CD25 | PC61 | PerCP/Cy5.5 | BD Biosciences |
| CD27 | LG.3A10 | PE | BD Biosciences |
| CD3e | 145-2C11 | PE/Cy5 | BD Biosciences |
| CD3e | 145-2C11 | FITC | BD Biosciences |
| CD3e | 145-2C11 | PB | BD Biosciences |
| CD4 | GK1.5 | FITC | BD Biosciences |
| CD45.1 | A20 | AlexaFluor700 | BioLegend |
| CD45.2 | 104 | BV650 | BioLegend |
| CD49b | DX5 | PE/Cy7 | eBioscience |
| CD69 | H1.2F3 | BV421 | BioLegend |
| DNAM-1 (CD226) | TX42.1 | BV510 | BioLegend |
| FLT3 (CD135) | A2F10.2 | BV421 | BioLegend |
| Gr1 (Ly6G/C) | RB6-8C5 | APC | BD Biosciences |
| Gr1 (Ly6G/C) | RB6-8C5 | FITC | BioLegend |
| IFNγ | XMG1.2 | BV605 | BioLegend |
| IgG2aκ | C1.18.4 | Purified | NordicBioSite |
| KLRG1 | 2F1/KLRG1 | BV421 | Biolegend |
| NK1.1 | PK136 | BV605 | BioLegend |
| NK1.1 | PK136 | Purified | BD Biosciences |
| NK1.1 | PK136 | Purified | NordicBioSite |
| NKp46 (CD335) | 29A1.4 | AF647 | BD Biosciences |
| Sca-1 | E13-161-7 | BV510 | BD Biosciences |
| Ter119 | Ter119 | PE; FITC | BD Pharmingen |

Supplementary Table II. List of Taqman probes (Applied Biosystems) used for Fluidigm gene expression analysis.

| Target gene | Taqman probe ID |
|---|---|
| *Akt* | Mm01331626_m1 |
| *B2m* | Mm00437762_m1 |
| *Bad* | Mm00432042_m1 |
| *Bcl2* | Mm00477631_m1 |
| *Bcl2l11* | Mm00437796_m1 |
| *Blimp1* | Mm00476128_m1 |
| *Cd160* | Mm00444461_m1 |
| *Cd2* | Mm00488928_m1 |
| *Cd27* | Mm01185212_g1 |
| *Cd94* | Mm00495182_m1 |
| *Cd96* | Mm00453394_m1 |
| *Cdkn1a* | Mm04205640_g1 |
| *Ccr2* | Mm00438270_m1 |
| *Ccr5* | Mm01216171_m1 |
| *Cdkn2b* | Mm00483241_m1 |
| *Cish* | Mm01230623_g1 |
| *Cxcr1* | Mm00731329_s1 |
| *Cxcr3* | Mm00438259_m1 |
| *Cxcr5* | Mm00432086_m1 |
| *Dap10* | Mm01172975_m1 |
| *Elf1* | Mm00468217_m1 |
| *Elf4* | Mm01321797_m1 |
| *Eomes* | Mm01351985_m1 |
| *Ets1* | Mm01175819_m1 |
| *Ezh2* | Mm00468464_m1 |
| *Fasl* | Mm00438864_m1 |
| *Foxo1* | Mm00490672_m1 |
| *Fyn* | Mm00433373_m1 |
| *Gata2* | Mm00492301_m1 |
| *Gata3* | Mm00484683_m1 |
| *Gata4* | Mm00484689_m1 |
| *Gzmb* | Mm00442834_m1 |
| *Hes1* | Mm01342805_m1 |
| *Hoxa9* | Mm00439364_m1 |
| *Hprt* | Mm00446968_m1 |
| *Id2* | Mm00711781_m1 |
| *Id3* | Mm00492575_m1 |
| *Inpp5d* | Mm00494987_m1 |
| *Irf1* | Mm01288580_m1 |
| *Ikzf1* | Mm00456421_m1 |
| *Ikzf3* | Mm01306721_m1 |
| *Il2ra* | Mm00434261_m1 |

| | |
|---|---|
| *Il2rb* | Mm00434268_m1 |
| *Il2rg* | Mm00442885_m1 |
| *Irf2* | Mm00515206_m1 |
| *Itgam* | Mm00434455_m1 |
| *Itgb7* | Mm00442916_m1 |
| *Jak1* | Mm00600614_m1 |
| *Jak3* | Mm00439973_m1 |
| *Jun* | Mm00495062_s1 |
| *Klf12* | Mm00516098_m1 |
| *Klf2* | Mm01244979_g1 |
| *Klra1* | Mm01183337_m1 |
| *Klra3* | Mm01702813_m1 |
| *Klra4* | Mm03647546_uH |
| *Klra6* | Mm00776306_mH |
| *Klra7* | Mm01183384_m1 |
| *Klrg1* | Mm00516879_m1 |
| *Klrk1* | Mm01183329_m1 |
| *mtor* | Mm00444968_m1 |
| *Ncr1* | Mm01337324_g1 |
| *Nfil3* | Mm00600292_s1 |
| *Mcl1* | Mm01257351_g1 |
| *Mysm1* | Mm00805196_m1 |
| *Pdcd1* | Mm01285676_m1 |
| *Pdk1* | Mm00554300_m1 |
| *Plcg1* | Mm01247293_m1 |
| *Prdm1* | Mm00476128_m1 |
| *Prf1* | Mm00812512_m1 |
| *Pten* | Mm00477208_m1 |
| *Ptpn6* | Mm00469153_m1 |
| *Ptpn11* | Mm00448434_m1 |
| *Rela* | Mm00501346_m1 |
| *Rora* | Mm01173766_m1 |
| *Rorc* | Mm01261022_m1 |
| *Runx1* | Mm01213405_m1 |
| *Runx3* | Mm00490666_m1 |
| *S1pr5* | Mm02620565_s1 |
| *Sell* | Mm00441291_m1 |
| *Sh2d1a* | Mm01316997_m1 |
| *Sh2d1b* | Mm00468982_m1 |
| *Smad2* | Mm00487530_m1 |
| *Socs3* | Mm00545913_s1 |
| *Stat5a* | Mm03053818_s1 |
| *Stat5b* | Mm00839889_m1 |
| *Spi1* | Mm00488140_m1 |
| *Tbx21* | Mm00450960_m1 |
| *Tcf3* | Mm01175588_m1 |

| | |
|---|---|
| *Tcf7* | Mm00493445_m1 |
| *Tigit* | Mm03807522_m1 |
| *Tnfsf10* | Mm00437174_m1 |
| *Tox* | Mm00455231_m1 |
| *Tyrobp* | Mm00449152_m1 |
| *Zap70* | Mm00494255_m1 |
| *Zbtb16* | Mm01176868_m1 |
| *Zeb2* | Mm00497193_m1 |
| *Zfp105* | Mm00494304_m1 |
